# TMPRSS6 cleaves KCNE1 and causes arrhythmias in iron overload disease

**DOI:** 10.1101/2024.09.04.611322

**Authors:** Stefan Peischard, Philipp Kastl, Gunnar Goerges, Julian A. Schreiber, Arie O. Verkerk, Ronald Wilders, Paul Disse, Isabelle Hornung, Ursula Klingmüller, Andrea U. Steinbicker, Martina Rauner, Maja Vujić Spasić, Frank Rosenbauer, Sven Meuth, Thomas Budde, Per A. Pedersen, Thomas A.Q. Jepps, Thomas Jespersen, Nathalie Strutz-Seebohm, Guiscard Seebohm

**Affiliations:** Institute for Genetics of Heart Diseases (IfGH), Department of Cardiovascular Medicine, University Hospital Münster, D-48149 Münster, Germany; Department of Systems Biology of Signal Transduction, German Cancer Research Center (DKFZ), D-69120 Heidelberg, Germany; DFG-research group FerrOs (FOR 5146), Germany; University of Münster, Institute for Pharmaceutical and Medicinal Chemistry, D-48149 Münster, Germany; Department of Medical Biology, Amsterdam Cardiovascular Sciences, Amsterdam UMC, University of Amsterdam, Amsterdam, Netherlands; DFG-Graduate School Chemical Biology of Ion Channels (Chembion), D-48149 Münster, Germany; Department of Anesthesiology, Intensive Care Medicine and Pain Therapy, University Hospital Frankfurt, Goethe University, D-60596 Frankfurt, Germany; Faculty of Medicine III and Center for Healthy Aging, Dresden University of Technology, D-01307 Dresden, Germany; Institute of Comparative Molecular Endocrinology, University of Ulm, Ulm 89081, Germany; Institute of Molecular Tumor Biology, University of Münster, 48149 Münster, Germany; Department of Neurology, University Hospital Düsseldorf, Medical Faculty, Heinrich Heine University, D 40225 Düsseldorf, Germany; Institute of Physiology I, Westfälische Wilhelms-Universität, Robert-Koch-Str. 27a, D-48149 Münster, Germany; Department of Biology, August Krogh Building, University of Copenhagen, Copenhagen, Denmark; Faculty of Health and Medical Sciences, Department of Biomedical Sciences, University of Copenhagen, Copenhagen, Denmark

## Abstract

Iron storage disease is associated with cardiovascular manifestations, including various forms of cardiac arrhythmias of unknown origin. In this study, cardiac arrhythmias associated with iron overload were investigated in human iPSC-derived cardiomyocytes (hiPSC-CM) and hiPSC-derived sinus node-like pacemaker cells. Among other effects, iron overload leads to an increase in the plasma membrane-anchored protease TMPRSS6. TMPRSS6 cleaves the auxiliary subunit KCNE1 N-terminally and thus modulates the function of both the I_Ks_ (KCNQ1/KCNE1 current) and the I_f_ (HCN4/KCNE1) ion channels. Furthermore, TMPRSS6 induces a reduction of electric field potential (EFP) count and increased duration in hiPSC-derived ventricular-like cells and in hiPSC-derived pacemaker-like cells. In accordance with these *in vitro* generated results, TMPRSS6-mediated interactions show pro-arrhythmic effects *in silico*. Therefore, the TMPRSS6 - KCNE1-KCNQ1 and TMPRSS6 - KCNE1-HCN4 cascades may represent new clinically relevant pro-arrhythmic mechanisms in iron overload diseases.

## Introduction

In iron storage disease (haemochromatosis, siderosis), the body is excessively overloaded with iron. There are two types of haemochromatosis, the primary and the secondary form. While primary haemochromatosis is a hereditary disease induced by genetic alterations of iron transporters, secondary haemochromatosis can be acquired as a result of certain anaemia subtypes, liver disease, thalassaemia or excessive blood transfusions ^1,2^. Symptoms include fatigue, exercise intolerance, fluid retention and dyspnoea. The excess iron can accumulate in organs and tissues causing gradual damage. While the liver is the main site of iron deposition in such patients, cardiac complications are the main cause of morbidity and mortality in haemochromatosis. Chronic iron overload leads to progressive cardiac dysfunction including heart failure and arrhythmia. In addition, haemochromatosis-related iron overload causes systemic oxidative stress, endothelial dysfunction and inflammation, all of which contribute to the development of atherosclerosis and an increased risk of cardiovascular events.

A genetic test following the diagnosis of haemochromatosis is positive in about three quarters of patients before symptoms develop ^3^. Early detection of iron storage disease in patients is urgently needed to initiate appropriate clinical treatment and prevent the progression of cardiac damage. Therefore, genetic testing is indicated as soon as clinical signs are detected.

The overall prevalence and risk of cardiovascular manifestations in patients with haemochromatosis at the population level have recently been investigated ^4,5^. Dilated cardiomyopathy (DCM) is the main cardiovascular manifestation of haemochromatosis. DCM is characterised by an enlargement of the heart and thus a disturbed geometry, which causes a reduced pump function ^6^. In addition, iron deposition in the myocardium triggers oxidative stress, inflammation, cell damage and fibrosis, which further reduce the heart’s contractility. Beside congestive heart failure and pulmonary hypertension, various forms of cardiac arrhythmias are frequent, with symptoms usually manifesting at higher ages ^4^. Iron overload prolongs the QT interval and reduces pace frequency, both being key elements for arrhytmogenesis in humans^7^. In a retrospective cohort analysis based on data from the National Inpatient Sample, Udani *et al*. and Jackson *et al*. ^4,5^ reported that 11.7% and 16.0% of haemochromatosis patients were diagnosed with cardiac arrhythmias, respectively, with an increasing prevalence in older patients. Male patients and non-Hispanic whites were more frequently affected compared to women and Hispanics. In women, symptoms of haemochromatosis, including severe arrhythmias, may not appear until later in life due to routine blood loss through the menstrual cycle. The haemochromatosis related cardiac arrhythmias include serious ventricular arrhythmias and heart block associated with higher in-hospital mortality, longer hospital stays and higher hospitalisation costs, which places a financial burden on healthcare systems ^4,8^. Population-level arrhythmias include supraventricular arrhythmias (10.6%), conduction of disturbances (2%), ventricular arrhythmias (0.8%) and cardiac arrest (0.4%) ^5^. The complex alterations in the electrical activity of the heart increase the risk of thromboembolic events and sudden cardiac death in those affected. Iron deposits in the conduction system cannot fully explain the complex electrical phenotype of the patients. The myocellular mechanisms of iron handling require further investigation.

Although iron is an essential nutrient, excess iron can be toxic to cardiac cells. The uptake of iron by mammalian cells is mediated by transferrin (Tf), which can coordinate two Fe^3+^ ions and binds to the transferrin receptor (TfR) on the plasma membrane of the cell. The Fe-Tf-TfR complex is endocytosed by vascular endothelial cells and cycles in the cytoplasm until it has reached a target within the cell. Post-transcriptional regulation through the interaction of the iron response proteins 1 and 2 (IRP1 and IRP2, respectively) with the iron response elements (IREs) located in the mRNAs of the iron metabolism genes represents an important feedback loop in cellular iron metabolism. Proteins, such as ferritin (for intracellular iron storage), the transferrin receptor, the erythroid aminolevulinic acid synthase, transmembrane serine protease 6 (TMPRSS6) and the mitochondrial aconitase, are regulated in this way. The IRPs bind in an iron-dependent manner to the IREs in the mRNAs of the regulated genes. The IRP interaction with the IREs changes the expression of the respective regulated genes. The induced altered gene expression of key proteins adapts the uptake, utilisation or storage of intracellular iron to the respective iron environment. In addition to iron content, the interaction of IRPs with IREs is sensitive to the redox state or the degree of oxidative stress in the cell, providing an important link between iron metabolism and the state of oxidative stress ^9^. Thus, post-transcriptional regulation of IRPs with IREs could regulate protein expression, which could affect the normal electrical function of the heart and lead to pro-arrhythmic effects.

Iron export from the cell represents a second pathway for cellular iron, namely transmembrane transport via ferroportin, the only known iron exporter on the cell surface. The secreted, hepatic peptide hepcidin negatively regulates cellular iron export by promoting the degradation of ferroportin ^10^ and thus adapting iron uptake to the body’s iron demand ^11^. Hepcidin is encoded by the HAMP gene and is itself negatively regulated by TMPRSS6 (encoded by the TMPRSS6 gene) ^12,13^. TMPRSS6 is a type II transmembrane serine proteinase that binds to arginines and cleaves proteins in the vicinity of serine residues ^14,15^.

Here, we show that iron elevation increases rhythmic heterogeneity in hiPSC cardiomyocytes (hiPSC-CM) with male genotype, which is a general pro-arrhythmic mechanism. To elucidate this effect, we analyzed protein expression altered by iron in hiPSC-CM. In agreement with previous reports, we find increased expression of the exoprotease TMPRSS6 upon iron overload *in vitro* ^16^. We identify the accessory ion channel subunit KCNE1 as a proteolytic target of TMPRSS6 and show that the functional interactions of both the cardiac pacemaker channel HCN4 and the delayed rectifier channel KCNQ1 with KCNE1 are altered by TMPRSS6 through cleavage of the extracellular amino terminus of KCNE1. Finally, TMPRSS6 modulates the electric activity of hiPSC-derived pacemaker cells and cardiomyocytes via KCNE1 modulation, thereby increasing the risk of complex cardiac arrhythmias.

## Results

### Iron overload and TMPRSS6 expression lead to cardiac arrhythmia

In recent years, the technique to transdifferentiate human iPS cells into different cardiac cell types has resulted in a robust source of specialized cardiac cells with similar characteristics such as ventricular-, atrial-, pacemaker- and conduction system-like phenotypes ^17–20^. Human iPS cardiomyocytes (ventricular-like type) were exposed to increasing Fe^2+^ levels for 3 days and rhythmic activities were analyzed by extracellular field potential (EFP) analysis. Even low Fe^2+^ concentrations lead to increasing rhythmic heterogeneity (Figure 1A). Rhythmic heterogeneity is a recognized pro-arrhythmic factor, which makes hiPSC-CM suitable for the analysis of iron-induced pro-arrhythmic mechanisms. The electric field potentials EFP/min count constantly decreased with increasing Fe^2+^ exposition (Figure 1B) and the average EFP duration increased (Figure 1C). Treatment of hiPSC-CM with the catalytic domain of the TMPRSS6 protease caused rhythmic heterogeneity similar to iron exposition (Figure 2A). In line with the EFP data in Figure 1B and 1C the incubation with TMPRSS6 decreased EFP/min (Figure 2B, left) and increased EFP duration (Figure 2B, right). In the liver, TMPRSS6 is upregulated following iron exposure ^10^. This type II transmembrane serine proteinase is embedded in the plasma membrane and could therefore be a potential modulator of plasma membrane-embedded ion channels involved in the proper rhythmicity of cardiomyocytes. To analyze whether iron exposure increases TMPRSS6 protein expression in hiPSC-CM, hiPSC-CM were incubated with 20 µM Fe^2+^ for 72 h and then stained via immunofluorescence against membrane-bound TMPRSS6. Indeed, Fe^2+^ incubation resulted in significant up-regulation of TMPRSS6 (Figure 2C). TMPRSS6 overexpression reduced the abundance of KCNE1, the regulatory β-subunit of the KCNQ1/KCNE1 complex. Using viral transfer of TMPRSS6 in mice, TMPRSS6 was expressed under a hepatocyte-specific promoter. Livers were subsequently obtained and then analyzed “for KCNE1 abundance via Western blot. TMPRSS6- overexpressing hepatocytes were found to contain significantly less KCNE1 than untreated cells (Figure 2D).

**Figure 1:**
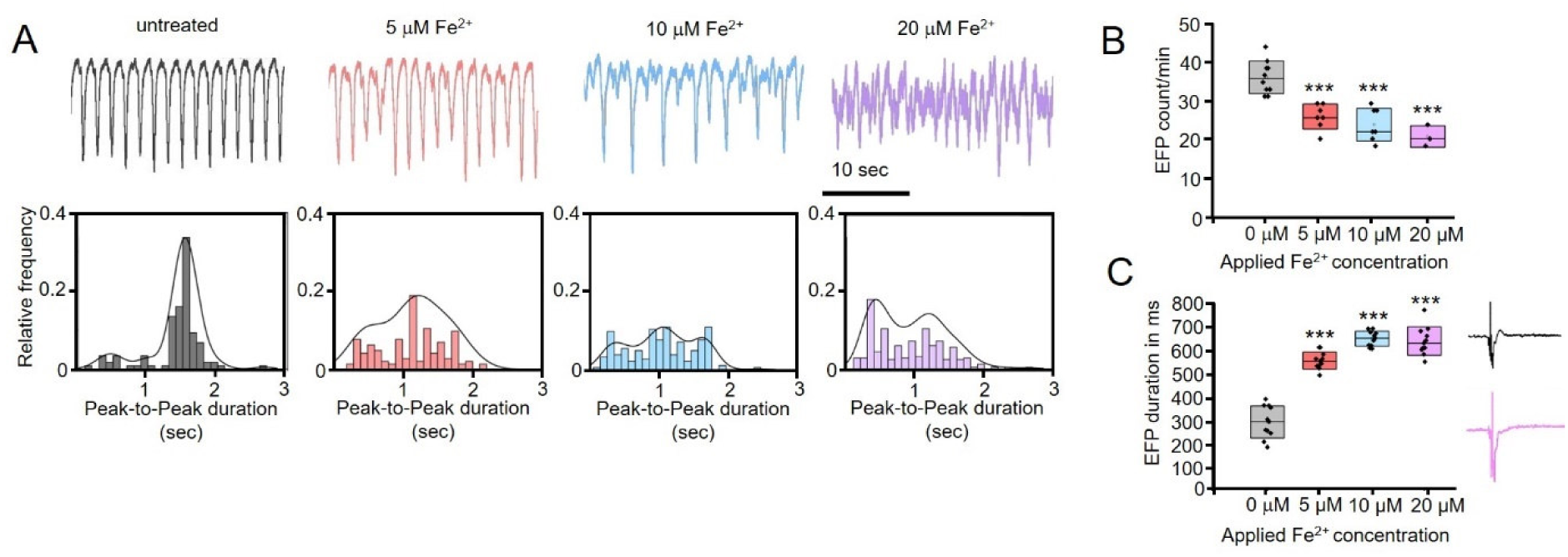
Identifying a new iron-dependent mechanism causing arrhythmicity in hiPSC-derived cardiomyocytes. **A;** Ventricular-like hiPSC-derived cardiomyocytes were incubated with rising concentrations of Fe^2+^ in the extracellular solution. The cell contractions were measured on the CardioExcyte 96 (CE96) and analyzed. Fe^2+^ incubation leads to contractile dysfunction already at a low concentration of 5 µM. Histograms in the lower panel include a Kernel Smooth fit to visualize the changes in contractility. **B;** The incubation of hiPSC-derived cardiomyocytes with Fe^2+^ reduces the production of electric field potentials (EFP) with rising iron concentrations. 0 μM (n=12), 5 μM and 10 μM (n=9), 20 μM (n=3) **C;** Elevated iron levels lead to the elongation of the EFP duration in hiPSC-derived cardiomyocytes 0 μM (n=10), 5 μM and 10 μM (n=7), 20 μM (n=3). \*\*\**p* ≤ 0.001.

**Figure 2:**
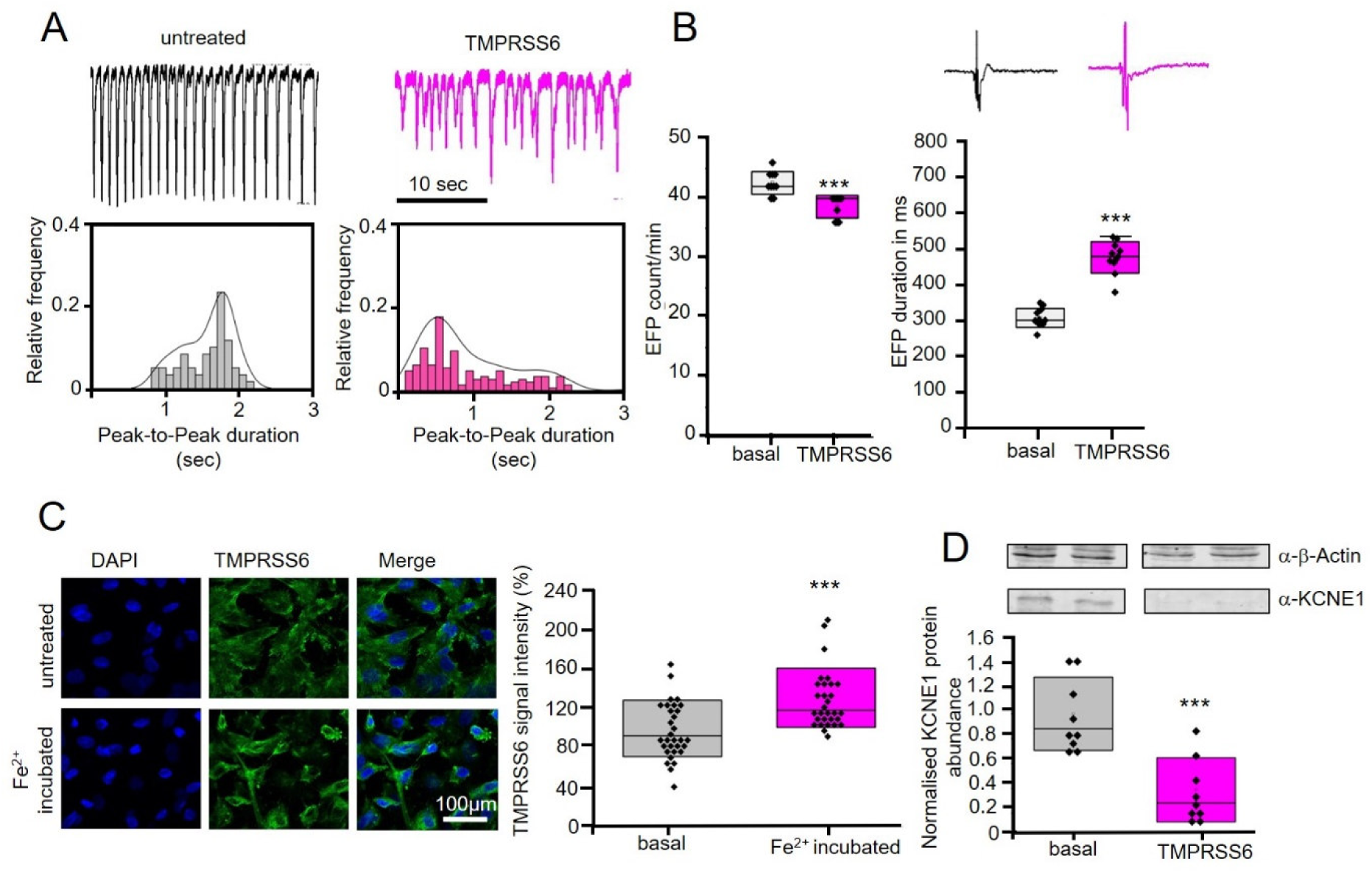
TMPRSS6 overabundance induces arrhythmia and is dependent on iron overload. **A;** HiPSC-derived cardiomyocyte contractions were measured on the CE96 under basal conditions (’untreated’) and under the influence of added TMPRSS6 to the medium. TMPRSS6 leads to similar contractile dysfunctions as seen in iron-incubated hiPSC-derived cardiomyocytes. Histograms in the lower panel include a Kernel Smooth fit to visualize the changes in contractility **B;** TMPRSS6 incubation of hiPSC-derived cardiomyocytes reduces EFP count (left, (n=10)) and elongates EFP duration (right, (n=11)) in hiPSC-derived cardiomyocytes. \*\*\**p* ≤ 0.001. **C;** Iron incubation leads to upregulation of TMPRSS6 in hiPSC-derived cardiomyocytes, control (n=31) and Fe^2+^ incubated (n=30). **D;** KCNE1 abundance is reduced by TMPRSS6 in mouse hepatocytes (n=9). \*\*\**p* ≤ 0.01.

### KCNQ1 currents are modulated by TMPRSS6

One of the most important determinants of the QT interval in humans is the cardiac K^+^ channel KCNQ1/KCNE1^21^. Decreased KCNQ1/KCNE1 activity leads to QT prolongation, which was observed in hiPSC-CM under iron exposure following incubation with TMPRSS6 (Figure 1C, 2B). Interestingly, this channel complex is sensitive to proteolysis by caspases ^22^ and by BACE ^23^. Two-electrode voltage clamp (TEVC) measurement of KCNQ1, KCNQ1 + KCNE1, KCNQ1 + TMPRSS6 and KCNQ1 + KCNE1 + TMPRSS6 showed slight decrease in KCNQ1 current when KCNE1 and TMPRS6 were co-expressed. The conductance was also shifted towards more positive potentials in oocytes co-expressing KCNQ1 + KCNE1 and TMPRSS6 (Figure 3A, B; Table 1). To test the effects of the TMPRSS6 proteolytic domain (PD) on human hiPSC-CM, we used hiPSC-CM with an endogenous KCNQ1 knockout, which allow doxycycline induction of KCNQ1 expression ^24^. The effects of TMPRSS6 proteolytc domain TMPRSS6-PD were tested in an isogenic approach in the absence or presence of the KCNQ1 channel α-subunit (Figure 3C-E). TMPRSS6-PD only affected the QT equivalent interval in the presence of KCNQ1 (Figure 3C). The EFP duration was increased in hiPSC-CM+KCNQ1+TMPRSS6 showing also recovery after washout (Figure 3D, top). The contraction amplitude was decreased in hiPSC-CM+KCNQ1+TMPRSS6 showing recovery after washout (Figure 3D, middle). The 50% contraction increase was not affected in any condition (Figure 3D, bottom). The rhythmic homogeneity was analyzed as well. HiPSC-CM-KCNQ1 were not affected by TMPRSS6 while hiPSC-CM+KCNQ1+TMPRSS6 showed an increase in rhythmic heterogeneity, which is in line with previous experiments (Figure 3E).

**Figure 3:**
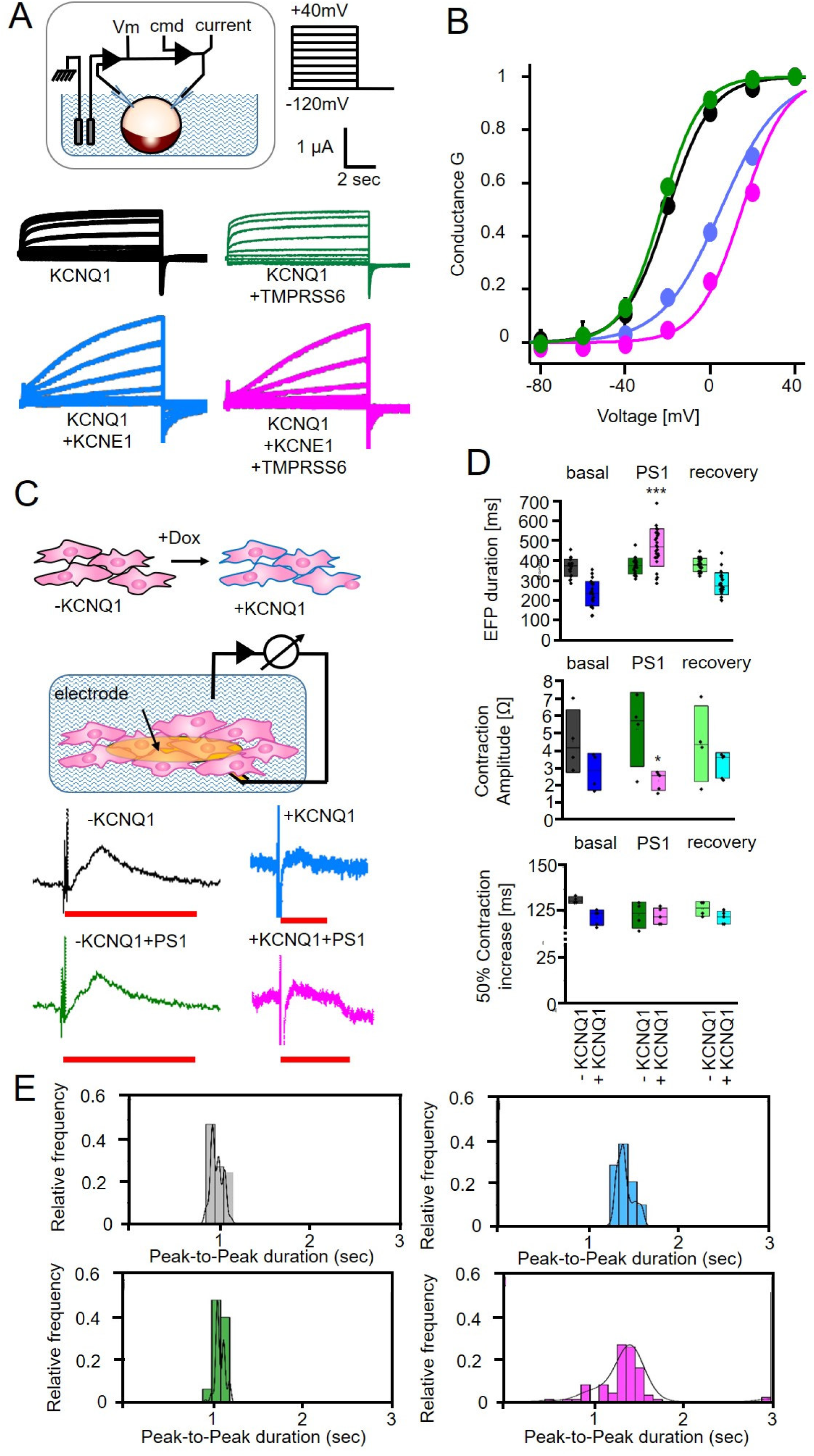
Identifying KCNQ1/KCNE1 as possible target for TMPRSS6. **A;** Two-Electrode-Voltage-Clamp of *Xenopus laevis* oocytes expressing KCNQ1 or KCNQ1 + KCNE1 measured under basal conditions and under co-expression of TMPRSS6. TMPRSS6 leads to slight decrease in KCNQ1 current when co-expressed with KCNE1. **B;** Voltage dependency of KCNQ1, KCNQ1 + KCNE1, KCNQ1 + TMPRSS6 and KCNQ1 + KCNE1 + TMPRSS6 co-expressing oocytes. Currents were normalized to +40 mV. TMPRSS6 leads to a shift in KCNQ1 excitability when co-expressed with KCNE1. **C;** CardioExcyte 96 measurement of hiPSC-derived cardiomyocytes of the cell line SFS.2-KCNQ1-GFP with endogenic KCNQ1 knockout and doxycycline-inducible KCNQ1 expression. TMPRSS6 addition (PS1) has no effect on QT interval (red bars) in KCNQ1KO cells while QT interval is elongated with PS1 when KCNQ1 is expressed. **D;** Analysis of EFP duration and contractility of SFS.2-KCNQ1-GFP cardiomyocytes. KCNQ1KO-CM did not show any effect of PS1 on EFP duration (n=20), contraction amplitude (n=4) and 50% increase time (n=4), while KCNQ1-expressing CM showed EFP elongation (n=25) and slight decrease of contraction amplitude (n=5). The 50% contraction increase time was unchanged (n=5). **E;** Analysis of SFS.1-KCNQ1-GFP contractility in response to PS1 addition. KCNQ1KO-CM did not respond to PS1 treatment while KCNQ1-expressing CM responded with arrhythmicity to PS1 addition. Histograms include a Kernel Smooth fit to visualize the changes in contractility \**p* ≤ 0.05; \*\*\**p* ≤ 0.001.

**Table 1:**
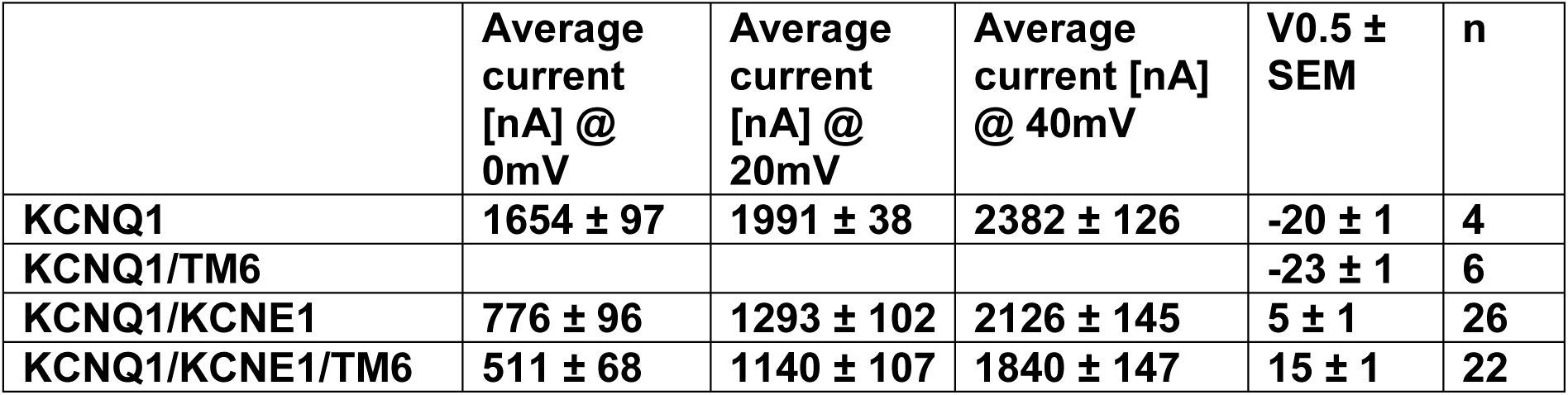
Results of TEVC experiments of KCNQ1, KCNQ1/KCNE1 and KCNQ1/KCNE1/TMPRSS6 co-expressing oocytes.

### KCNE1 is cleaved by TMPRSS6

The fact that TMPRSS6-PD acts on cardiomyocytes via the KCNQ1/KCNE1 channel complex suggests that the channel may be cleaved via PD. The type II transmembrane serine proteinase TMPRSS6 is embedded in the plasma membrane and cleaves proteins at RS and KS motifs ^25^. Inspection of the KCNQ1 α-subunit did not reveal a potential cleavage site. However, the KCNE1 β-subunit contains a potential arginine cleavage motif (^32^RRSPRS^37^) near the outer membrane (Figure 4A). Mass spectrometry (MS) can be used to verify the cleavage of synthetic peptides by the purified TMPRSS-PD (Figure 4B). Several synthetic peptides representing the extracellular domain (ECD) of KCNE1 were used to estimate the cleavage products generated upon application of TMPRSS-PD (Figure 4C). The results confirm that TMPRSS-PD can hydrolyze the extracellular domain (ECD) of KCNE1 at arginines *in vitro* (Figure 4D). To verify the cleavage of KCNE1 ECD and the dependence of cleavage on arginines, a novel fluorescence-based *in vitro* assay was developed (Figure 5A). A KCNE1 construct with external N-terminal Myc+pHmScarlet (red fluorophore) and internal C-terminal EGFP was constructed. Transiently transfected HEK 293T cells with robust plasma membrane expression could be FACS-sorted via red pHmScarlet fluorescence, as expression of the construct in intracellular vesicles is easily overlooked by the low vesicular pH, which reduces pHmScarlet emission ^26^ (Figure 5B). Extracellular pHmScarlet is cleaved together with the extracellular domain of KCNE1, if TMPRSS6 is present and pHmScarlet signal intensity is reduced (Figure 5B). One set was co-transfected with an empty pcDNA3 vector as a mock control. A second set of HEK 293T cells was analogously transfected with an additional px-Flag construct to express TMPRSS6. To normalize the effect of co-transfected TMPRSS6 with the constructs pHmScarlet-KCNE1-GFP (WT or R32A, R33A, R36A), a mock control was also prepared and analyzed. FACS analysis of pHmScarlet-KCNE1(mutant)-GFP-transfected HEK 293T cells showed that the basal number of pHmScarlet-positive cells was comparable for all transfected constructs (WT, R32A, R33A, R36A), as shown in Supplementary Figure 1. By transfecting KCNQ1WT with a mock control, the effect of co-transfection on the resulting pHmScarlet fluorescence signal was determined and used to normalize the results of samples transfected with KCNE1 variants + TMPRSS6. Analysis of samples co-transfected with TMPRSS6 clearly showed that KCNE1WT was cleaved by TMPRSS6, as evidenced by a 15.58% reduction in the number of pHmScarlet-positive cells. This effect was reduced to the level of the mock control in the R32A, R33A and R36A mutants, indicating the importance of these three arginines for TMPRSS6 proteolysis *in vivo* (Figure 5C). In conclusion, the KCNE1-ECD is a TMPRSS6 substrate for proteolysis *in vitro* and *in vivo*.

**Figure 4:**
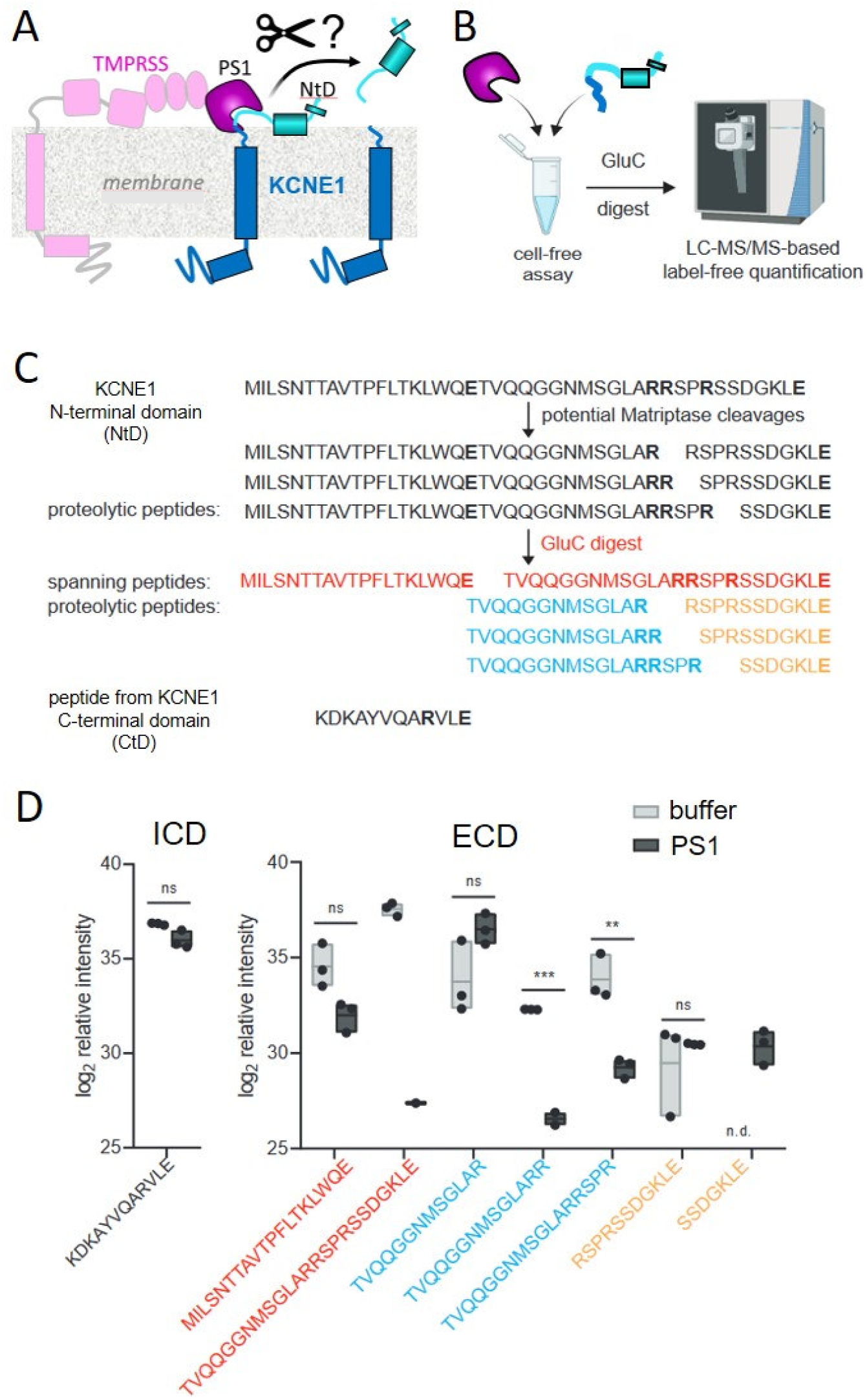
Confirmation of Matriptase-mediated cleavage of the KCNE1 extracellular domain (ECD) at arginines by mass spectrometry. **A;** Concept of Matriptase-mediated cleavage of KCNE1. **B**; Mass spectrometry-based experimental approach to confirm Matriptase-mediated cleavage of the KCNE1-ECD with label-free quantification (LFQ). **C;** Amino acid sequence of the ECD of KCNE1 and theoretically generated peptides after tryptic digest with GluC (blue) as well as potential Matriptase cleavage sites (red arginines) and the resulting proteolytic peptides. **D**; Log2 LFQ intensities of spanning and proteolytic peptides after incubation of the KCNE1-ECD with buffer control or recombinant Matriptase (n=3). Dots are single values, bars represent the means ± SEM. \*\**p* ≤ 0.01; \*\*\**p* ≤ 0.001.

**Figure 5:**
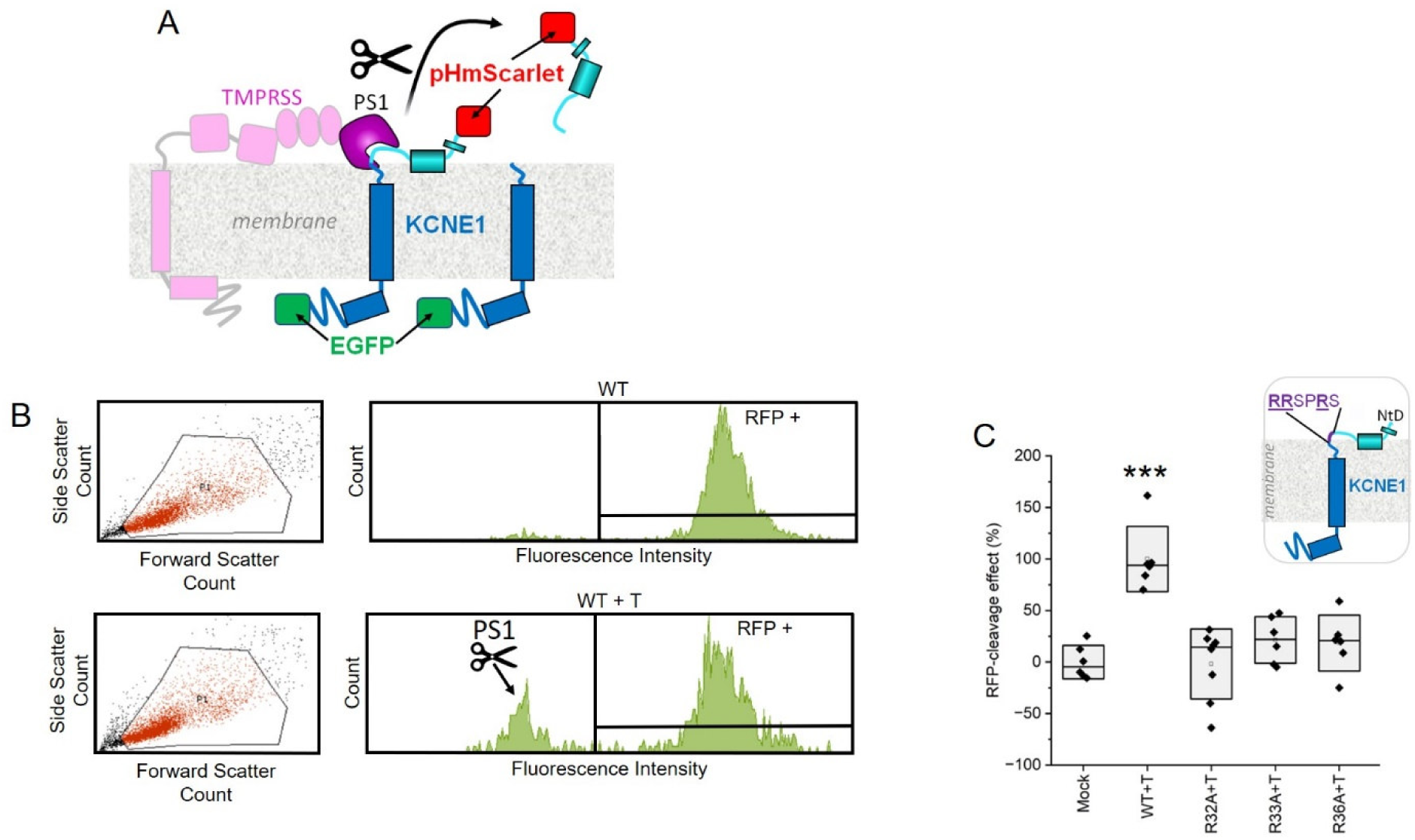
TMPRSS6 cleaves KCNE1-protein in heterologous expression depending on membrane proximal arginines. **A;** Concept of pHmScarlet cleavage of transfected KCNE1-pHmScarlet constructs co-expressed with TMPRSS6. Extracellular pHmScarlet is cleaved by TMPRSS6 if a TMPRSS6 recognition site is located at the extracellular compartment of KCNE1. The resulting red fluorescence intensity can be analyzed and quantified via FACS. **B;** Exemplary FACS analysis of HEK cells transfected with KCNE1WT-pHmScarlet ± TMPRSS6 (T). KCNE1WT-pHmSarlet + T shows an increase in the RFP low intensity cell fraction proving the responsiveness of the generated KCNE1-pHmScarlet construct to TMPRSS6. **C;** KCNE1 mutants of the KCNE1-pHmScarlet construct were generated at a predicted TMPRSS6 recognition site (RRSPRS) in which the inherited arginines (R) were substituted with alanines (A) resulting in the constructs KCNE1(R32A, R33A and R36A). Co-expression with the KCNE1-pHmScarlet constructs revealed significant effect on TMPRSS6 on the KCNE1WT but not on the mutants KCNE1(R32A; R33A and R36A) (n=6).; \*\**p* ≤ 0.01; \*\*\**p* ≤ 0.001.

### KCNE1 is cleaved by TMPRSS6 at a specific cleavage site

To combine the experimental data with structural analyses of the peptide/protease complex, we constructed a TMPRSS6-KCNE1 protein complex model (Figure 6A, 6B). The position of the KCNE1 key region ^31^ARRSPRSG^38^ with respect to the substrate cleft of the TMPRSS6 protease was determined by analogous positioning to the peptidomimetic benzothiazole that was previously co-crystallized with the homologous peptidic matriptase-1 (TMPRSS2) (PDB ID 6N4T). Molecular dynamics (MD) simulations were then performed in a membrane. The simulations revealed that the KCNE1 31-36 region and TMPRSS6 form a stable complex that is anchored to the membrane. The KCNE1 residues ^41^KLEALYVLMVLGFFGFFTLGIMLSYIR^67^ form a transmembrane α-helix, as shown previously ^27^. The N-terminal region of TMPRSS6 also contains a transmembrane α-helix (^46^PLFVLLALLVLASAGVLLWYFLGYK^70^). These two regions anchor the TMPRSS6-KCNE1 complex to the membrane (Figure 6B). Thus, the TMPRSS6-CUB1 - CUB2 - LDL receptor homologous region and the protease domain are aligned parallel to the membrane. For equilibration, the TMPRSS6-KCNE1 complex was simulated for 30 ns embedded in a membrane surrounded by a water shell leading to a constant fluctuating root mean square deviation (RMSD; Figure 6C). Afterwards, three independent production runs (sim 1-3) were conducted for an additional 270 ns leading to a total simulation time of 3 × 300 ns for the complex. Within this simulation time, KCNE1_31-36_ adopted a stable conformation within the catalytic peptidase S1 domain (PS1) without disrupting the active side of the domain (Figure 6D). Moreover, several H bond and ionic interactions stabilize KCNE1_31-36_ in close proximity to the catalytic triad that is located at S753 (nucleophile), H608 (base, H bond acceptor) and D659 (deprotonation of H608) (Figure 6E, 6F) ^25^. Analyses of the interaction endurance between KCNE1 and TMPRRS6 show that interactions clustering around the proteolytic residue S753 (H608, D659, D747, V772, S773, W774, G775, G777) persist for more than 70% of the mean production simulation time (Figure 6G). In particular, interactions between KCNE1 R36 and TRPMSS6 D747 as well as G777 via H bond and / or ionic interaction were detected in at least 98% of the analyzed simulation snapshots positioning R36 in close proximity to S753 and increasing the chance of nucleophilic attack and subsequent proteolytic cleavage (Figure 6G, 6H).

**Figure 6:**
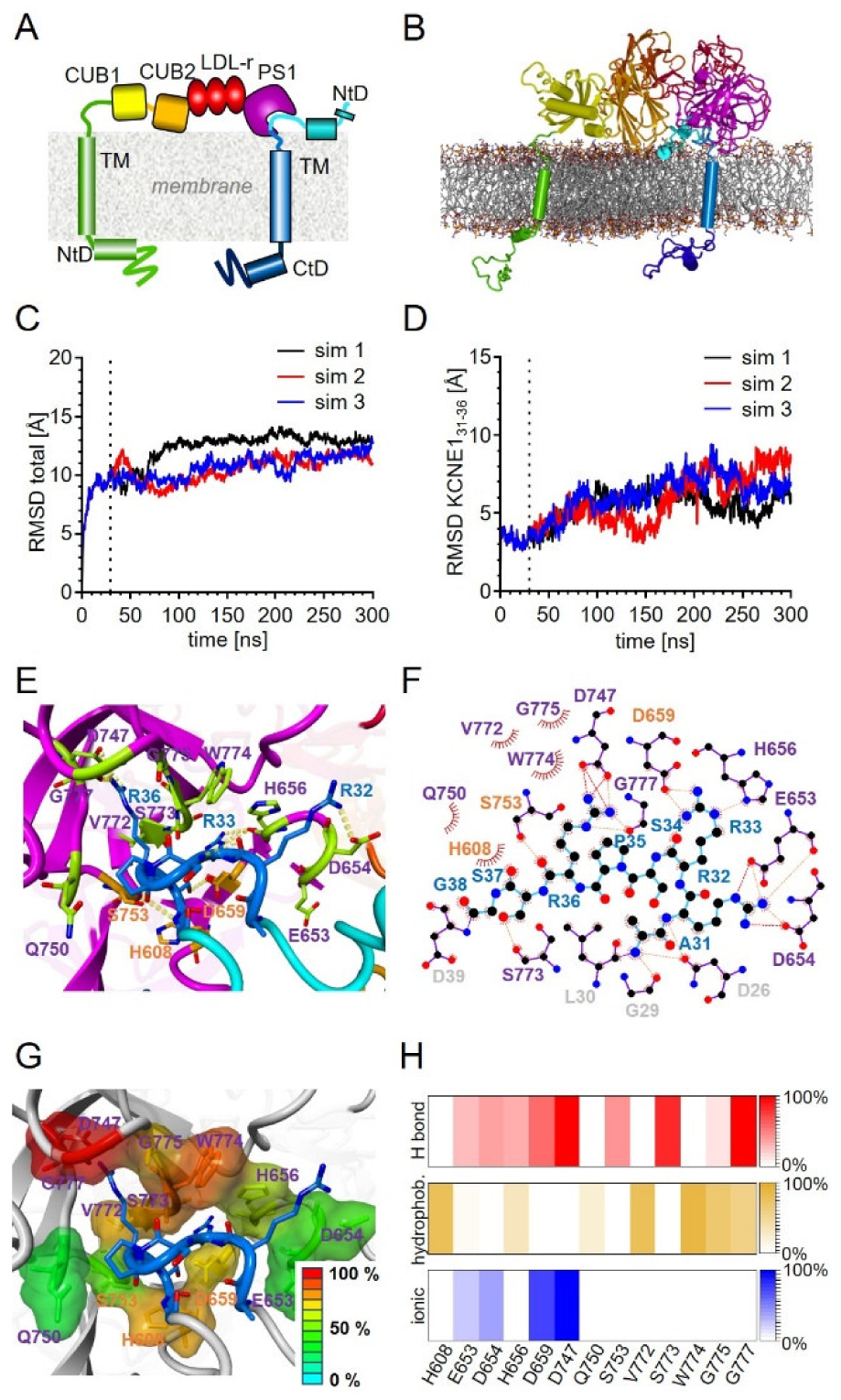
KCNE1 forms a stable complex with TMPRSS6 *in silico*, enabling proteolytic cleavage. **A, B;** The TMPRSS6-KCNE1 model complex was constructed as described in the Methods section. The arrangement of structural motifs is shown as cartoon (A) and in simplified structural form (B). TMPRSS6 consists of an intracellular N-terminal domain (NtD, residues 1-209, green) with a transmembrane α-helix (TM, residues 46-69), the CUB1 domain CUB1, residues 210-336 (yellow), a second CUB2 domain (CUB2, residues 337-452, orange), LDL receptor class 1-3 domains (LDL-r, residues 453-567, red) and the catalytic peptidase S1 domain (PS1, residues 568-802, magenta). KCNE1 contains an extracellular N-terminal domain (NtD, residues 1-40, light blue), an α-helical transmembrane domain (TM, residues 41-67, blue) and an intracellular C-terminal domain (CtD, residues 68-129, dark blue). **C;** Total root-mean-square deviation (RMSD) [Å] of TMPRSS6 / KCNE1 complex for equilibration and production simulations (sim 1-3). Start of production runs is indicated by black dotted line. **D;** RMSD [Å] of KCNE1 residues A31-G38 for equilibration and production simulations. Black dotted line indicates the start of production runs. **E;** Depiction of KCNE1_31-38_ (dark blue) bound to the PS1 domain (magenta) of TMPRSS6. Interacting amino acids of TMPRSS6 are highlighted in green or orange (catalytic triad H608, D659 and S753). H bonds between KCNE1_31-38_ and PS1 domain are indicated by yellow dotted line. **F;** Depiction of H bond (orange line) and ionic (red line) interactions between KCNE1_31-38_ (blue) and PS1 (purple, orange). Further interacting residues of KCNE1 are indicated in gray. **G;** Structural depiction of interacting PS1 residues colored by interaction persistence over the production simulation time ranging from 0% (blue) to 100% (red). Residue colors are derived from mean interaction percentage of three independent simulations. **H;** Persistence of H bond, hydrophobic and ionic interactions over the production simulation time represented as mean from three independent simulations ranging from 0% (white) to 100% (dark color) of the simulation time.

### HCN4 currents are affected by TMPRSS6

KCNE1 also affects the function of the human sinus node pacemaker channel HCN4 ^28^. Indeed, we confirm that KCNE1 increases HCN4 currents in a heterologous expression system (Figure 7A, 7B). However, coexpression of HCN4/KCNE1 channels with TMPRSS6 partially diminishes the KCNE1-stimulation on HCN4 currents (Figure 7A, 7B; Table 2). Noteworthy, TMPRSS6 coexpression exerts no effect on HCN4 without KCNE1, underscoring that TMPRSS6 effects depend on the presence of KCNE1 (Figure 7A, 7B).

**Figure 7:**
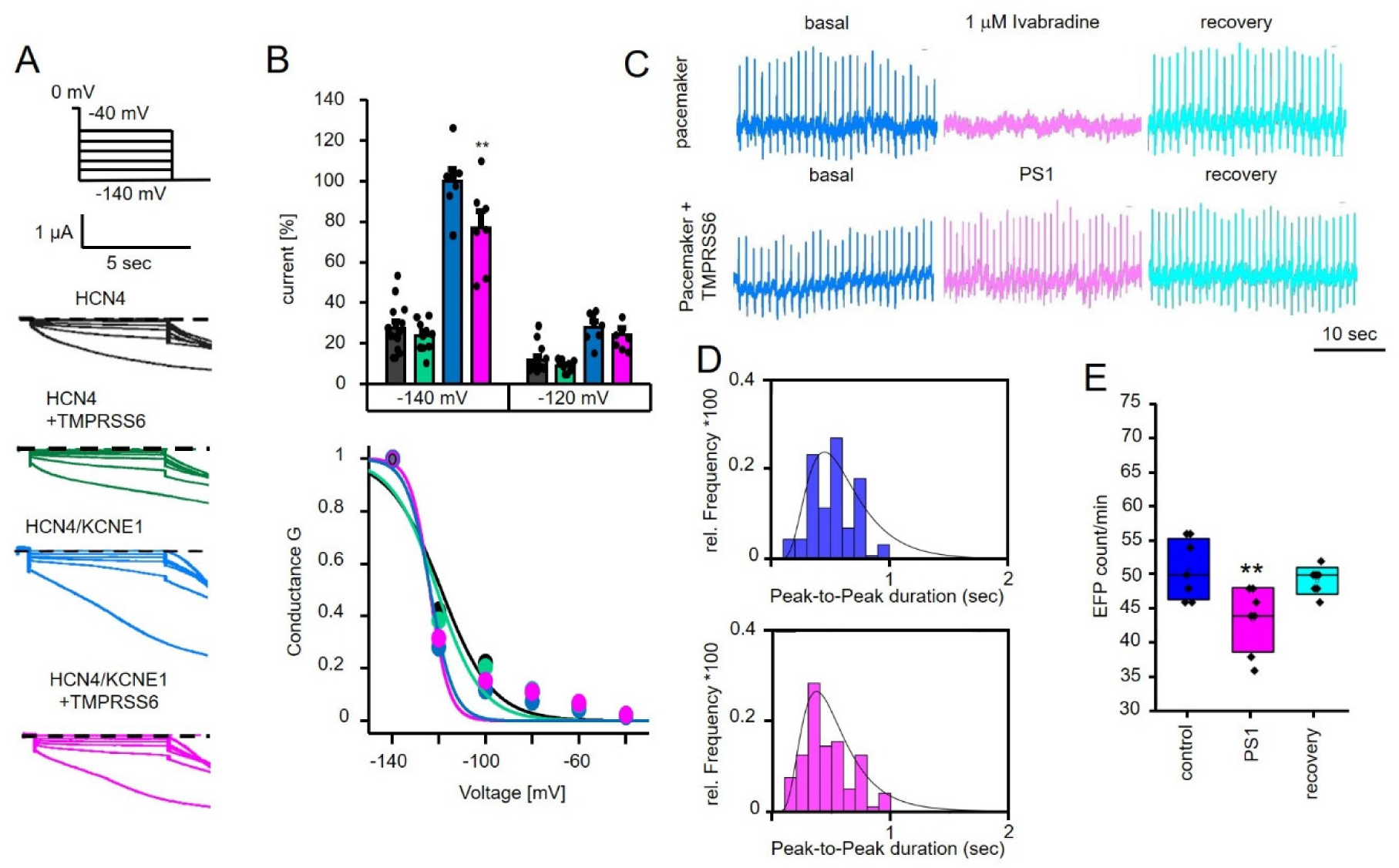
Identifying the cardiac pacemaker channel HCN4 as second target for TMPRSS6-induced KCNE1 cleavage. **A;** Two-Electrode-Voltage-Clamp of *Xenopus laevis* oocytes expressing HCN4 or HCN4 + KCNE1 measured under basal conditions and under co-expression of TMPRSS6. TMPRSS6 leads to decrease in HCN4 current when co-expressed with KCNE1. **B;** Statistical analysis of HCN4 currents measured in TEVC. The bars show the conditions HCN4 (black), HCN4 + TMPRSS6 (green), HCN4 + KCNE1 (blue) and HCN4 + KCNE1 + TMPRSS6 (magenta) at −140 mV and −120 mV (top figure) (n=10). Conductance effect of TMPRSS6 induced KCNE1 cleavage on HCN4 was evaluated with no observable effect (bottom figure). **C;** HiPSC-derived cardiomyocytes with ventricular or sinus-nodal phenotype were measured with the CE96. HiPSC-derived sinus-nodal cells (pacemaker cells) react highly sensitive to 1 µM Ivabradine, proving that hiPSC-derived pacemaker cells are highly HCN4 dependent. TMPRSS6-PS1 treatment of hiPSC-derived pacemaker cells has no macroscopic effect on pacemaker cell activity. **D;** HiPSC-derived pacemaker cells were measured on the CE96 under basal conditions (blue) and treated with TMPRSS6-PS1 (magenta). Slight effect on contractility was observed after TMPRSS6-PS1 treatment. Histograms include a Kernel Smooth fit to visualize the changes in contractility **E;** The count of EFP/min was evaluated under basal conditions and under the influence of TMPRSS6-PS1 addition in hiPSC-derived pacemaker cells. TMPRSS6-PS1 leads to significant reduction of the EFP count (n=7). \*\**p* ≤ 0.01.

**Table 2:** Results of TEVC experiments with HCN4, HCN4/KCNE1 and HCN4/KCNE1/TMPRSS6 co-expressing oocytes.

|  | Average current [%] @ -120mV | Average current [%] @ -140mV | V0.5 ± SEM | n |
| --- | --- | --- | --- | --- |
| HCN4 | 12 ± 2 | 28 ± 3 | -119 ± 1 | 14 |
| HCN4/TM6 |  |  |  |  |
| HCN4/KCNE1 | 28 ± 3 | 100 ± 6 | -124 ± 1 | 7 |
| HCN4/KCNE1/TM6 | 24 ± 3 | 77 ± 8 | -124 ± 1 | 7 |

HCN4 channels trigger the action potentials in the human sinus node and hiPSC pacemaker cells ^29^. Inhibition of HCN4 in hiPSC pacemaker cells by ivabradine inhibits beating in the cells suggesting that pacemaking in the cells critically depends on HCN4 channel function (Figure 7C). Application of the TMPRSS6-catalytic domain PS1 slightly increased rhythmic heterogeneity in hiPSC pacemaker cells consistent with a reduction in HCN4 function (Figure 7D). Thus, TMPRSS6-PS1 reduces HCN4/KCNE1 function and reduces EFP count/min in hiPSC pacemaker cells (Figure 7E).

### Mathematical modelling identifies the importance of KCNE1 cleavage on KCNQ1 and HCN4 activity

We carried out computer simulations to quantify the effects of the TMPRSS6-induced decrease in I_Ks_ and the TMPRSS6-induced decrease in I_f_ on the electrical activity of single human sinus node pacemaker cells. First, we assessed the effects of the TMPRSS6-induced decrease in I_Ks_ *per se*. As illustrated in the leftmost panels of suppl. Figure 8, the amplitude of I_Ks_ is almost halved upon the TMPRSS6-induced shift in the voltage dependence of I_Ks_. This decrease in outward current results in a slight increase in the rate of diastolic depolarization and, consequently, a slight increase in pacemaker rate. Next, we assessed the effects of the TMPRSS6-induced decrease in I_f_ *per se*. As illustrated in the middle panels of suppl. Figure 8, the amplitude of I_f_ is slightly reduced upon the TMPRSS6-induced decrease in the fully-activated conductance of I_f_. The decrease in inward current during diastolic depolarization results in a slight decrease in diastolic depolarization rate and pacemaker rate. Finally, we assessed the combined effect of the TMPRSS6-induced decrease in I_Ks_ and I_f_. As illustrated in the rightmost panels of suppl. Figure 8, the combined effect is rather small, with a decrease in pacemaker rate <3%. Similar results were obtained at other levels of autonomic tone (Supplementary Figures 4-6) and in the presence of a passive atrial load (Supplemental Figures 7-9). These simulation results are in line with the experimental observation of Figure 7C that TMPRSS6-PS1 treatment of hiPSC-derived pacemaker cells has no macroscopic effect on pacemaker cell activity. Human ventricular cardiomyocytes do not functionally express I_f_, but they do express I_Ks_. Therefore, we also carried out computer simulations to assess the effects of the TMPRSS6-induced decrease in I_Ks_ in human ventricular tissue. To this end, we constructed a linear strand of 400 cells (Figure 8A) that we stimulated at one end at rates of 60 and 180 beats/min. At a rate of 60 beats/min, the generated action potential (AP) successfully travels through the strand and is identical in shape for cells #100 and #300 of the strand, both under control conditions and in the simulated presence of TMPRSS6, which has an AP prolonging effect (Figure 8B). This also holds for APs that are elicited at a rate of 180 beats/min (Figure 8C). However, essentially different results are obtained with the functionally more demanding branched strand of Figure 8D. At a rate of 60 beats/min, the APs are still successfully conducted throughout the strand and are again identical in shape for cells #100 and #300 of the strand, both under control conditions and in the simulated presence of TMPRSS6 (Figure 8E). However, at a rate of 180 beats/min and in the simulated presence of TMPRSS6, every second AP in the unbranched part of the strand is not completely full-blown, resulting in an alternating long-short AP pattern (Figure 8F, left, green traces). The short AP is not successfully conducted to the branched part of the strand and this partial block thus results in a 2:1 conduction pattern in the strand (Figure 8F, right, green traces). Of note, this pro-arrhythmic partial block was not observed under control conditions (Figure 8F, blue traces).

**Figure 8:**
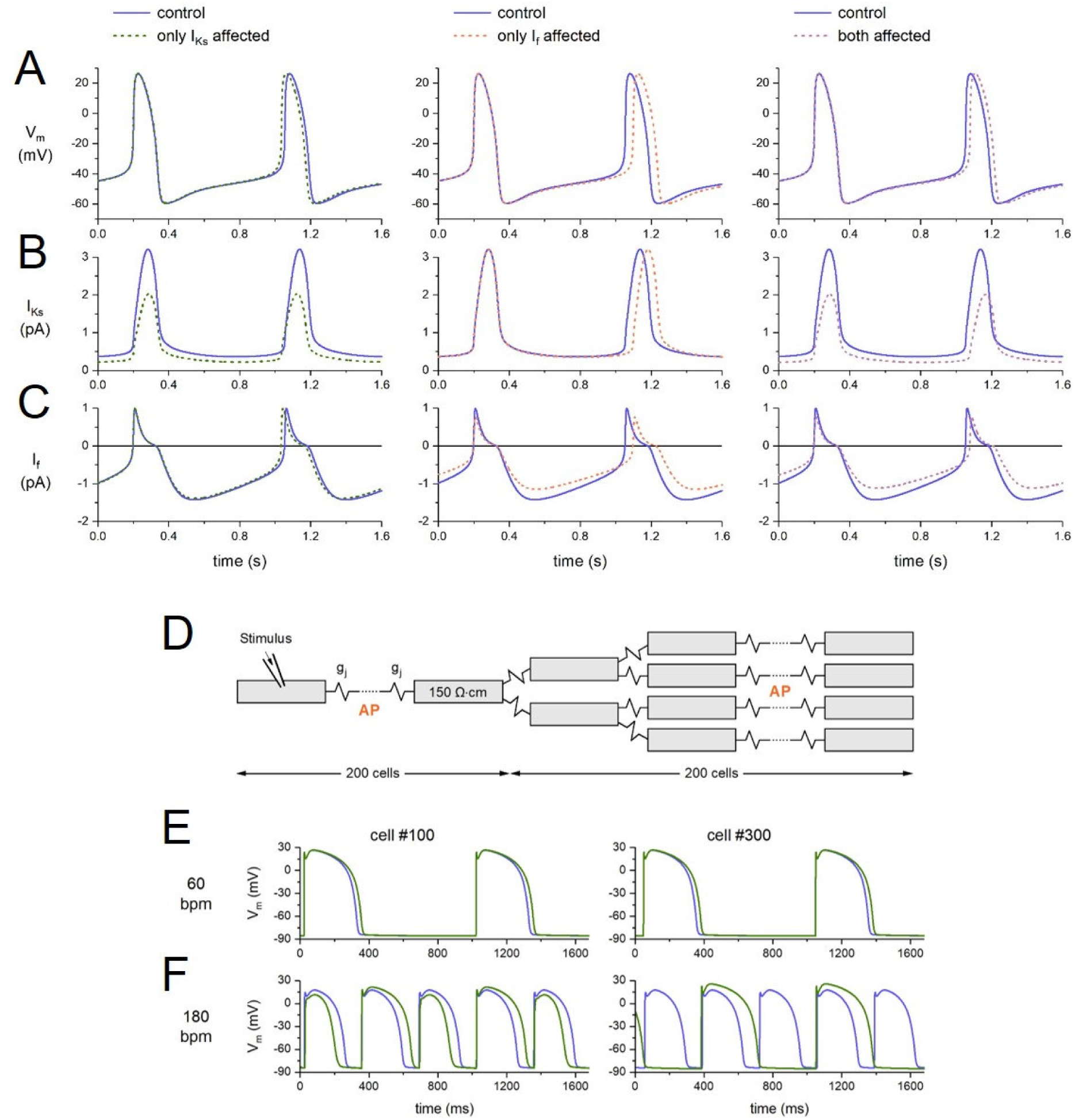
*In silico* effects of TMPRSS6 on the electrical activity of a single human sinus node cell and effects of TMPRSS6 on the electrical activity in a simulated linear strand of human left ventricular cardiomyocytes under control conditions (blue traces) and during the TMPRSS6-induced reduction in I_Ks_ (green traces). **A;** Membrane potential (V_m_), **B;** slow delayed rectifier potassium current (I_Ks_), and **C;** hyperpolarization-activated ‘funny’ current (I_f_) when TMPRSS6 affects I_Ks_ only (left), I_f_ only (middle), or both I_Ks_ and I_f_ (right). **D;** Geometry of the branched strand of 400 cells. The positions of cells #100 and #300 in the strand is labeled with ‘AP’. **E;** V_m_ of cells #100 (left) and #300 (right) at a stimulus rate of 60 beats/min. **F;** V_m_ of cells #100 (left) and #300 (right) at a stimulus rate of 180 beats/min.

## Discussion

A comprehensive understanding of the pathogenesis of cardiovascular manifestations in haemochromatosis is required for the development of new appropriate therapeutic approaches. As novel therapeutic targets for a systemic disease need to be identified, a multidisciplinary approach involving knowledge from cardiology, hematology and other fields is required to obtain a complete picture of iron overload disease. The basic mechanisms of the cardiac arrhythmias associated with iron overload are still largely unexplored and cellular or organoid systems may prove suitable to model the complex disease. Here, we utilize current human iPSC techniques to generate ventricular cardiomyocyte-like cells and sinus node-like pacemaker cells. Increased iron concentration in the medium leads to arrhythmias in ventricular-like cardiomyocytes in 3D cultures (Figure 1). Polymorphic arrhythmias are common in the iron overload disease haemochromatosis. Patients’ arrhythmias include, in particular, supraventricular arrhythmias, conduction disturbances, and potentially fatal ventricular arrhythmias and heart block ^5,8^. The phenotype of cardiac arrhythmias in ventricular-like cardiomyocytes in 3D cultures appears to be complex and corresponds to the variable and complex phenotype of cardiac arrhythmias in patients. Thus, the hiPSC-CM model represents a suitable model for arrhythmias in iron overload.

The response to an increase in extracellular iron has been extensively studied in mouse liver models. These studies have shown that TMPRSS6 expression is modulated by iron status ^30,31^. Iron regulatory proteins 1 and 2 (IRP1 and IRP2) post-transcriptionally control the expression of several mRNAs encoding proteins involved in iron metabolism. In particular, IRP1 is an important regulator of erythropoiesis and iron uptake by controlling mRNA translation of hypoxia-inducible factor 2α (HIF2α). Cellular studies in Hep3B cells have shown that TMPRSS6 is one of the key players in iron metabolism, which is upregulated by HIF-1α and HIF-2α. This upregulation of TMPRSS6 leads to a decrease in membrane hemojuvelin and thus to a decrease in hepcidin production in mouse liver ^32^. Here we show that elevated iron levels increase TMPRSS6 in hiPSC-CM (Figure 2). It is possible that a similar iron-regulated IRP1-HIF-2α pathway as in the liver also regulates cardiac TMPRSS6 expression. Further studies are required to clarify this point.

TMPRSS6 is a membrane-bound arginine-serine protease (peptidase S1-type) that cleaves membrane-bound hemojuvelin in the extracellular region ^33^. It is possible that such protease activity can cleave ion channel-related proteins and thus modulate the function of cardiac ion channels to cause cardiac arrhythmias. This hypothesis would involve proteolysis of target auxiliary subunits and was tested by applying the purified catalytic domain of TMPRSS6 to chambered cardiomyocytes in 3D cultures (Figure 2). Indeed, application of the catalytic domain of TMPRSS6 induced arrhythmias is similar to the application of iron, suggesting that iron-induced upregulation of TMPRSS6 is a crucial mechanism for triggering arrhythmias.

One of the most important determinants of cardiac action potential duration in humans is the cardiac KCNQ1/KCNE1 K^+^ channel ^34,35^. Interestingly, this channel complex is sensitive to proteolysis by caspases ^22^ and by BACE ^23^. Heterologous co-expression of the KCNQ1/KCNE1 channel with full-length TMPRSS6 identifies TMPRSS6 as a novel regulator of the potassium channel. TMPRSS6 causes a rightward shift in the voltage dependence of KCNQ1/KCNE1, which decreases the overall current density (Figure 3A, 3B). Fast activating KCNQ1 α-subunit is insensitive to co-expression with TMPRSS6, suggesting that the TMPRSS6 sensitivity of the channel complex is mediated by the KCNE1 β-subunit (Figure 3A, 3B). The reduction in native cardiac K^+^ current I_Ks_, which is conducted by KCNQ1/KCNE1 channels, is clinically associated with drug-induced long QT syndrome and inherited long QT syndromes 1 and 5, which induce ECG abnormalities and cause arrhythmias as well as sudden cardiac death (summarized by Sanguinetti and Seebohm ^21^). In particular, the ventricular action potential is prolonged due to the decrease in or complete absence of the repolarizing I_Ks_, which can be observed in the surface ECG as a prolonged QT interval. The ECG aberrations due to the absence of KCNQ1/KCNE1 can be well studied in hiPSC-CM in a tet-on-KCNQ1 system in a KCNQ1KO background^24^. The effect of TMPRSS6-PD on the QT-equivalent interval depends on the presence of the KCNQ1 channel α-subunit (Figure 3C, 3D), emphasizing that TMPRSS6-PD acts via the KCNQ1/KCNE1 channel complex.

Since TMPRSS6 acts on the KCNQ1 channel complex via KCNE1, we investigated whether TMPRSS6 cleaves KCNE1. The potential TMPRSS6 cleavage site in the KCNE1 region ^29^GLARRSPRSSDG^39^ was cleaved in a cell-free assay (Figure 4). In this assay, cleavage is critically dependent on the three inherited arginines. Although a cell-free assay can detect physical cleavage, it does not prove that this cleavage also *occurs in cells*. Therefore, a novel cell-based assay was developed. The attachment of fluorophores to the extracellular N-terminus and the intracellular C-terminus enables the differentiation between transfected cells with reduced extracellular fluorophore (Figure 5A, 5B). To favor KCNE1 inserted into the plasma membrane, the extracellular fluorophore is pH-sensitive (pHmScarlet is an RFP-like fluorophore) and the intracellular fluorescence of this potentially cleaved fluorophore is reduced due to acidification in the vesicles. Use of this assay demonstrated that TMPRSS6 cleaves a population of labeled KCNE1 and confirmed that the arginines R^32^, R^33^ and R^36^ are also required for cleavage in cells (Figure 5C). The correct positioning of the ^29^GLARRSPRSSDG^39^ motif can be proposed as the basis for targeted cleavage.

The structure of TMPRSS6 in complex with a peptidomimetic benzothiazole (PDB 6N4T) and the structure/dynamics of KCNE1 (PDB 2K21) were determined ^25,27^ and used to generate a TMPRSS6-KCNE1 structural model (see Methods). These modeling efforts demonstrate that the arginines R^32^, R^33^, and R^36^ are critical for the proper interaction of KCNE1 with the cleavage site in TMPRSS6 *in silico* and provide a structural explanation for the relevance observed in cell-free assays and in cells (Figures 3, 4, and 6).

The KCNE1 β-subunit has been proposed as a clinically relevant modulator of a second cardiac ion channel, the pacemaker channel HCN4 ^29,36,37^. This channel controls the electrical activity of the sinus node both *in vivo* and in 3D cultures of hiPSC-derived pacemaker cells ^29,38–40^. Loss-of-function of HCN4 by several mechanisms has been associated with hereditary sinus node dysfunction and atrial/ventricular tachyarrhythmias ^29,41–43^. Coexpression of HCN4/HCNE1 with TMPRSS6 but not HCN4 without KCNE1 showed decreased current density (Figure 7A, 7B) with heterologous expression. Such decrease is observed at −140 mV, but not at less negative potentials. The reduced current density of HCN4/KCNE1 currents by TMPRSS6 is consistent with reduced pacemaker function in 3D cultures of hiPSC-derived pacemaker cells (Figure 7D, 7E), suggesting that native HCN4 current I_f_ is modulated by TMPRSS6-cleaved KCNE1 in these cells.

Physiological diurnal variations in plasma iron levels have been reported ^44^ and diurnal variations in the QT interval and heart rate are well known as well. However, extensively increased serum iron has been associated with prolongation of the QT interval and reduced cardiac pace ^7^. Indeed, *Fabbri–Severi* model simulations suggest that TMPRSS6-induced reduced pace and reduced I_Ks_ compensate each other on the level of the action potential (suppl. Figure 8) and suggesting that these changes may contribute to cardiac diurnal adaptations. However, the human SA node cell simulations indicate that potentially pro-arrhythmic effects may be prominent under increased β–adrenergic tone implying sympathetic tone as relevant pro-arrhythmic factor (Figure 8; Suppl. Figure 6). Atrial load (passive load mimicking atrial load) or increased vagal tone seem to be less relevant for TMPRSS6 effects, as suggested by human SA node cell simulations in the *Fabbri–Severi* model (Suppl. Figures 4-9).

In summary, iron overload induces an increase in TMPRSS6, which in turn cleaves KCNE1, modulating both I_Ks_ (KCNQ1/KCNE1 current) and I_f_ (HCN4/KCNE1 current) via KCNE1 cleavage. TMPRSS6-mediated effects on cardiac currents were included in mathematical models and indeed show pro-arrhythmic effects *in silico* (Figure 8). Therefore, the iron-induced cascades TMPRSS6 - KCNE1-KCNQ1 and TMPRSS6 - KCNE1-HCN4 described here may represent relevant pro-arrhythmic mechanisms in iron overload diseases. These novel cascades open the way to new therapeutic treatment options and identify new antiarrhythmic drug targets in iron overload.

## Methods

### FACS analysis

HEK 293T cells were seeded into the wells of a 24-well plate at a density of 300,000 cells per well and allowed to grow overnight. The next day, the cells were transfected. Prior to transfection, the medium was changed to 300 µL HEK cell medium and then the transfection mix was prepared. The transfection mix for the transfection of one well consisted of 25 mL DMEM (D5671, Sigma Aldrich), 2 µg of each DNA to be transfected and PEI (3 µL/µg DNA). The prepared reaction mixture was allowed to sit for 20 minutes and then applied dropwise to the cells. The transfected cells were incubated at 37°C and 5% CO_2_ for 6 hours. After 6 hours of incubation, the medium was replaced with 1 mL of HEK cell medium. The next day, the HEK cell medium was removed and the cells were washed once with PBS. The cells were detached with 1 mL trypsin-EDTA solution (T3924-500ML, Merck) for 2 minutes. The trypsin reaction was stopped by adding 1 mL of HEK cell medium to the cells. The detached cells were transferred to a 2 mL Eppendorf tube and centrifuged at 200 × g for 3 minutes. The supernatant was removed and the cells were fixed with 4% PFA solution for 10 minutes. The fixed cells were centrifuged at 200 × g for 3 minutes. The supernatant was removed and the cells were washed with PBS. The cells were centrifuged again at 200 × g for 3 minutes. The supernatant was removed and the cells were resuspended in 0.5 mL PBS. The cell suspension was transferred to a FACS tube (352235, Corning) and analyzed by FACS. Cells were first sorted by green fluorescence to ensure that only positively transfected cells were analyzed and then analyzed by their red fluorescence intensity to quantify the effect of TMPRSS6. The threshold for a positive red fluorescence signal was determined by analyzing non-transfected HEK 293T cells.

### HEK cell culture

HEK 293T cells were seeded and cultured in HEK cell medium consisting of DMEM (D5671, Sigma Aldrich), 1× penicillin/streptomycin (P4333-100ML, Sigma Aldrich), 10% FBS superior (S0615, Merck) and 1× Non-Essential Amino Acids (M7145-100ML, Sigma Aldrich) in a T-25 flask (833910002, Sarstedt). When 90% confluence was reached, the medium was removed and the cells were washed once with PBS. A volume of 2 mL trypsin-EDTA (T3924-500ML, Merck) was added to the cells and incubated at 37°C for 2 minutes. The trypsin reaction was stopped by adding 5 mL of HEK cell medium to the cells. The cells were isolated and collected in a 15 mL Falcon tube. After centrifugation at 200 × g for 2 minutes, the resulting cell pellet was resuspended in 10 mL HEK cell medium. The cells were seeded at a ratio of 1:10 into a new T-25 flask. The remaining cells were used for the experiments.

### HEK cell transfection

HEK 293T cells were seeded into the wells of a 24-well plate (833922, Sarstedt, Germany) at a density of 300,000 cells/well in HEK cell medium. The cells were incubated overnight at 37°C and 5% CO_2_. The next day, the medium was replaced with 0.5 mL HEK cell medium. The transfection mixture was then prepared. 25 µL DMEM and 2 µg DNA were mixed in a 1.5 mL Eppendorf tube. A volume of 3 µL/µg DNA PEI (43896.01, VWR) was added to the transfection mixture. The mixture was incubated for 30 minutes at room temperature. The incubated transfection mixture was added dropwise to the respective 24 wells containing HEK cells. The transfected cells were incubated for 6 hours at 37°C and 5% CO_2_. After 6 hours, the medium was changed to 1 mL HEK cell medium. The next day, the cells were prepared for fluorescence analysis.

### Animals

The animal protocols were approved by the Regierungspräsidium Darmstadt (approval number FK/2028). Eight-week-old male C57BL/6 mice were injected intravenously with either 1×10^12^ particles of AAV2/8 expressing TMPRSS6-FLAG under the control of a liver-specific promoter (AAV8-TBG-mTMPRSS6.FLAG, Vector BioLabs) or PBS as control. Three weeks after virus administration, mice were euthanized under deep anesthesia and organs were harvested for further analysis.

### Immunoblotting

Liver samples were lysed in RIPA buffer supplemented with protease and phosphatase inhibitor cocktails (Sigma-Aldrich), and protein concentration was determined with the Pierce BCA Protein Assay Kit (Thermo Fisher Scientific). Equal amounts (50 µg) of isolated proteins were subjected to 16% SDS-PAGE under denaturing and reducing conditions (10% β-Mercaptoethanol), and samples were incubated for 5 min at 95°C prior to loading. Proteins were transferred to nitrocellulose membranes and incubated with primary antibodies directed against Kcne1 (PA5-106872, Thermo Fisher Scientific) and β-Actin (A5441, Sigma-Aldrich). Immunoblots were then incubated with appropriate secondary antibodies (926-68073 and 926-32212, Li-Cor). Imaging was performed using the Odyssey CLx Imaging System (Li-Cor).

### HiPSC culture

Human induced pluripotent stem cells from Sendai-Foreskin 1 (SFS.1/ KCNQ1-GFP) were cultured on 6-well plates coated with a 1:75 dilution of Matrigel®, and the culture medium was changed daily. KO-DMEM was used for the Matrigel® dilution supplemented with 5% KnockOut Serum Replacement (KSR) and 1 × penicillin/streptomycin/glutamine (PSG). The maintenance medium, FTDA medium, consisted of DMEM/F12, 1 × PSG, 1% CD lipid concentrate, 5 µg/mL ITS, 0.1% human serum albumin (HSA), 10 ng/mL FGF-2, 0.2 ng/mL TGF-β, 50 nM dorsomorphin and 5 ng/mL Activin A. Cells were divided every 4 days after reaching complete confluence. At division, cells were washed with phosphate-buffered saline (PBS), detached with 1 mL of Accutase solution supplemented with 10 µM Y-27632 ROCK inhibitor, and then incubated at 37°C for 10-12 min until complete detachment. Subsequently, 1 mL FTDA + 10 µM Y-27632 ROCK inhibitor was added to the cells and the cell suspension was transferred to a 15 mL Falcon tube. After centrifugation at 200 × g for 3 min, the supernatant was removed, and the cells were resuspended in 5 mL FTDA + 10 µM Y-27632 ROCK inhibitor. Approximately 650,000 cells were then seeded onto new Matrigel®-coated 6-well plates.

### Cardiogenic differentiation of hiPSC

Cells of the SFS.1/KCNQ1-GFP cell line were seeded on day 0 in differentiation medium in 24-well plates coated with 1:400 diluted Matrigel® at a density of 500,000 cells per well. The differentiation medium on day 0 contained KO-DMEM, 1 × ITS, 10 μM Y-27632, 1 × PSG, 5 ng/mL Activin A, 10 ng/mL FGF-2, 0.5-1 ng/mL BMP4 and 1 μM CHIR99021. After 24 hours, the medium was replaced with TS-ASC medium consisting of KO-DMEM, 1 × PSG, 1 × transferrin/selenium solution and 250 μM 2-phospho-ascorbate. Cells were treated daily with 2 mL/well of TS-ASC medium. On days 2 and 3 of differentiation, the WNT inhibitor C59 was added at a final concentration of 0.2 µM. Differentiation was continued until day 8, when the cells spontaneously beat.

For ventricle-like differentiation, beating cardiomyocytes were transferred to Matrigel®-coated plates with additional gelatin coating after 8 days. Cells were divided and cultured in KO-THAI medium consisting of KO-DMEM, 1 × PSG, 1 × ITS, 0.2% HSA, 250 μM phospho-ascorbate and 0.008% thioglycerol supplemented with 10 μM Y-27632. The cells were cultured in KO-THAI for 28 days to achieve a more mature ventricular-like phenotype ^17,29^.

For pacemaker-like differentiation, beating cardiomyocytes were transferred to Matrigel®-coated plates with gelatin coating after 8 days. Cells were divided into new wells at a ratio of 1:4 and cultured in KO-THAI medium containing 10 μM Y-27632. Twenty-four hours later, the medium was replaced with pacemaker differentiation medium consisting of KO-DMEM, 1 × PSG, 1 mM CaCl_2_, and 10% FBS. The cells were cultured in pacemaker differentiation medium for 28 days to achieve a pacemaker-like phenotype ^17,29^.

### CardioExcyte 96

HiPSC-derived cardiac-like cells were seeded on 0.1% gelatin-coated NSP-96 Type Standard Stim plates (# 201003, Nanion Technologies GmbH) at a density of 30,000 cells/well. The cells were seeded in their respective medium (KO-THAI for ventricular-like cells and Pacemaker medium for SAN-like cells) with an additional 10 µM Y-27632 and incubated at 37°C and 5% CO_2_ for 24 hours. The next day, the medium was replaced by the respective medium of the cells without Y-27632. The cells were left untreated until spontaneous contraction occurred, which was 5 days after seeding. The NSP-96 Type Standard Stim plate seeded with hiPSC-derived cardiac-like cells was inserted into the CardioExcyte 96 (Nanion Technologies GmbH) for measurement. Spontaneous cell activity (electric field potential generation EFP and contraction across the baseline impedance) was measured for 2 h every 15 min for 30 s at 37°C and 5% CO_2._ For the assessment of MMP-2 activity on the respective cell type, 10u MMP-2 (ALX-201-752-0500, ENZO) was added to the desired wells. Cell activity (electric field potential generation EFP and contraction across the base impedance) was monitored for a further hour. The medium was then changed in all wells and the cells were incubated at 37°C and 5% CO_2_ overnight (12 h) for recovery. Cell activity (generation of electric field potential EFP and contraction across baseline impedance) was monitored for one hour after recovery. Data analysis was performed using CardioExcyte Data Control Software (Nanion Technologies GmbH).

### Test for cleavage of the extracellular domain of KCNE1

The synthetic peptides were resuspended in 80% 0.1 M ammonium bicarbonate and 20% acetonitrile to a final concentration of 1 mM. For the matriptase cleavage assay, a peptide mixture containing 100 µM of total KCNE1-ECD and 33 µM each of three different peptides of KCNE1-ICD was mixed with either equal volumes of assay buffer (50 mM Tris, 50 mM NaCl, 0.01% (v/v) Tween® 20, pH 9.0) or 500 ng of recombinant human matriptase/ST14 catalytic domain (R&D Systems #3946-SEB-010) and incubated at 37°C for 2 hours. Treatment with buffer and matriptase was performed in triplicate.

### Sample preparation for mass spectrometry analysis

Peptides cleaved with matriptase and incubated with buffer were mixed with Sera-Mag carboxylate-modified magnetic beads (100 µg/µL) (Cytiva) at a ratio of 10:1 (beads:protein) and ethanol was added to a final concentration of 50%. The mixture was incubated for 15 minutes at room temperature and then washed three times with 80% ethanol on a magnetic rack. The peptide-bound beads were resuspended in 75 µL of 100 mM triethylammonium bicarbonate, pH 7.5, and digested with GluC (1:50 protease:peptide) for 16 h at 37 °C. The digested samples were separated from the beads on a magnetic rack, dried by vacuum centrifugation (45°C, 1300 rpm, RVC 2-18 CDplus CHRIST) and stored at −20 °C.

### Mass spectrometric measurement

Samples were analyzed using an Orbitrap Exploris 480 coupled to an UltiMate 3000 RSLCnano system (Thermo Fisher Scientific) equipped with a trapping column (precolumn; C18 PepMap 100, 5 µm, 300 µm i.d. x 5 mm, 100 Å). The samples were resuspended in loading buffer (0.1% formic acid, 2% acetonitrile in HPLC water) and the peptide content was determined using a Nanodrop DS-11+ spectrophotometer, (DeNovix), 600 ng loaded onto the trapping column (30 µL/min 0.5% TFA in HPLC water for 3 min) and eluted from an analytical column (75 mm × 30 cm, packed with Reprosil-Pur 120 C18-AQ, 1.9 µm resin, Dr. Maisch, Ammerbuch, Germany) for 85 min with a stepped gradient of 8-38% solvent B (0.1% formic acid, 80% acetonitrile in HPLC water; solvent A: 0.1% formic acid in HPLC water) at a flow rate of 300 nL/min with a final 5 min step of 98% solvent B. The mass spectrometer was operated in data-dependent mode (DDA) with automatic switching between MS and MS2. Full scan MS spectra in the Orbitrap were acquired with a resolution of 60,000 and a mass range of 350 to 1500 m/z. Tandem mass spectra were generated for up to 20 peptide precursors in the Orbitrap (isolation window 1.0 m/z) for fragmentation using high energy collision dissociation (HCD) at a normalized collision energy of 30% (Orbitrap Exploris 480) and a resolution of 15,000 (Orbitrap Exploris 480) with a target value of 100,000 charges after a maximum accumulation of 96 ms. The raw MS spectra were processed with MaxQuant (version 2.0.1.0) for peak detection and quantification. MS/MS spectra were searched using the Andromeda search engine against a Fasta file consisting of only bovine albumin (P02769), human ST14 (Q9Y5Y6) and human KCNE1 (P15382), allowing contamination detection and the detection of reversed versions of all sequences with the following search parameters: acetyl (protein N-term) and oxidation (M) as variable modifications. GluC was specified as the proteolytic enzyme with semi-specificity and no missed cleavages allowed. The mass accuracy of the precursor ions was determined by the time-dependent recalibration algorithm of MaxQuant. The maximum false discovery rate (FDR) for proteins and peptides was 0.01, and a minimum peptide length of six and maximum of 35 amino acids was required. Protein quantification was done using the label-free quantification algorithm implemented in MaxQuant. All other parameters were set to the default settings. All proteome data are deposited with the ProteomeXchange consortium via the partner repository PRIDE ^45^ with the dataset identifier PXD054634 with the username: and password: a5YRXwLDYBf1.

### Two-electrode voltage clamp (TEVC) in *Xenopus* laevis oocytes

cRNA synthesis, oocyte preparation and two-electrode voltage clamp were performed as previously described ^42,46,47^.

Specifically, cDNA was linearized and transcribed into cRNA using an *in vitro* transcription system (mMESSAGE mMACHINE T7 Transcription Kit, Invitrogen, Carlsbad, USA). Defolliculated oocytes (EcoCyte Bioscience, Dortmund, Germany) were injected with different combinations of HCN4 (5 ng), KCNQ1 (2.5 ng), KCNE1 (2.5 ng) and TMPRSS6 (5 ng) cRNA. After injection, oocytes were incubated in Barth’s solution (88 mM NaCl, 1 mM KCl, 0.4 mM CaCl_2_, 0.33 mM Ca(NO_3_)_2_, 0.6 mM MgSO_4_, 5 mM TRIS-HCl, 2.4 mM NaHCO_3_, supplemented with 80 mg/L theophylline, 63 mg/L benzylpenicillin, 40 mg/L streptomycin and 100 mg/L gentamycin) for 2 days. TEVC measurements were performed at a holding potential of −80 mV for KCNQ1 and −40 mV for HCN4-expressing oocytes using recording pipettes filled with 3 M KCl (resistance 0.5-1.5 MΩ) in ND96+ solution (96 mM NaCl, 4 mM KCl, 1.8 mM MgCl_2_, 0.1 mM CaCl_2_, 5 mM HEPES; adjusted to pH 7.4 with 1 M NaOH).

The pulse protocol shown in Figure 3A was used to examine voltage-dependent activation of KCNQ1 channels. In HCN4-expressing oocytes, pulse protocol 1 (Figure 7A) was used to determine the voltage-dependent activation of HCN4 channel currents, while a pulse protocol 2 was used (Figure 7B) to assess the voltage-dependent deactivation of HCN4 channel currents. Both pulse protocols were applied to the same oocyte. Data were recorded using GePulse (Dr. Michael Pusch, Genoa, Italy).

### Molecular modeling

Molecular modeling was performed with YASARA 23.9.29, LigPlot+ v2.2 and OriginPro 2019 for data analysis. ^48^ The KCNE1 structures of the protein studied by NMR in micelles (2K21.pdb) were adopted from Kang et al. ^26^ The AlphaFold model of TMPRSS6 and the crystal structure of matriptase-1 in complex with a peptidomimetic benzothiazole (6n4t.pdb) from Béliveau et al. ^25^ were used to build a consensus model using the Yasara homology modeling approach. The peptidomimetic benzothiazole were (6n4t.pdb) identifies the exact position of the key arginine that enables the positioning of target peptides to the arginine-directed serine protease TMPRSS6. This arginine corresponds to KCNE1-R^36^, which precedes the cleavage target S37. KCNE1-R36 was used to position the KCNE1 key region ^31^ARRSPRSG^38^ in the interaction region of the TMPRSS6 protease gap analogous to peptidomimetic benzothiazole (6n4t.pdb). The KCNE1 region was kept flexible while TMPRSS6 was fixed. Docking steps followed by energy minimization were performed until no significant changes in the docked KCNE1 peptide were observed. This TMPRSS6-KCNE1 complex was energy minimized using unchanged YASARA equilibration method and represented our starting model for further structural studies. For the MD simulation in a lipid membrane, the provided YASARA 23 macro md_runmembrane.mcr was used and the following parameters were adjusted: The membrane composition was set to 100% phosphatidyl ethanolamine (PEA), all polar head groups were substituted with 1-palmitoyl- and 2-oleoyl. Positioning of the TMPRSS6-KCNE1 model followed the standard Yasara unbiased procedure. The position was assumed to be predicted and was not changed manually. The simulation was conducted using the following settings: AMBER14 force field, simulation time steps of 2*2.5 fs, periodic simulation cell boundary, particle-mesh Ewald / Poisson-Boltzmann cutoff 8 Å, water model TIP3P (density 0.997 including 0.9 % NaCl, pH 7.4), 1 atm pressure and temperature of 298 K. All simulations were controlled by an NPT ensemble like previously described ^49^. Simulation time was set to 30 ns for equilibration followed by three independent production runs for 270 ns each. Root mean square deviation (RMSD) for the whole TMPRSS6 / KCNE1 complex as well as for KCNE1_31-36_ was monitored over the complete simulation time every 0.25 ns (equilibration + production). For interaction analyses, snapshots of the production runs were analyzed (3 x 1080 snapshots). For this purpose, the YASARA macro md_analyze.mcr was adapted defining KCNE1_31-36_ as a ligand. H bond, hydrophobic and ionic interactions were analyzed using predefined parameters from the md_analyze.mcr macro. Interactions were analyzed and counted for each snapshot and normalized to the number of production snapshots (1080 snapshots). Interacting residues were identified and analyzed with a cutoff of at least 20% interaction time between the residue and KCNE1 over the whole production simulation time.

### Action potential modeling

The electrical activity of a single human sinus node pacemaker cell was simulated using the comprehensive model of such a cell developed by Fabbri et al. ^50^ and known as the Fabbri–Severi model, with updated equations for the slow delayed rectifier potassium current (I_Ks_) ^51^. A passive atrial load was simulated by the addition of an outward current with a constant conductance and a reversal potential of −80 mV. The CellML code ^52^ of the Fabbri–Severi model, available from the CellML Model Repository ^53^ at https://models.cellml.org/e/568/ (accessed on July 30, 2024), was edited and run in version 0.9.31.1409 of the Windows-based Cellular Open Resource (COR) environment ^54^. All simulations were run for a period of 100 s to ascertain stable behavior. The data analyzed are from the final five seconds of this 100 s period.

The electrical activity in a linear strand of 400 human ventricular cardiomyocytes with or without branches was simulated using the Ten Tusscher et al. ^55^ human left ventricular cell model, as updated by Ten Tusscher and Panfilov ^56^, to describe individual cells in the strand. The myoplasmic resistivity was set to 150 Ω·cm ^57^ and the gap junctional conductance was set to 10 µS ^58^. The number of cells in the branched and unbranched strands was large enough to rule out any effects of the ‘sealed ends’ at both ends of the strand. Software was coded and compiled as a 32-bit Windows application using Intel Visual Fortran Composer XE 2013 and run on an Intel Core i7 processor-based workstation. For the numerical integration of differential equations we applied a simple and efficient Euler-type integration scheme with a 1 µs time step ^59^. All simulations were run for a period of 50 stimuli to ascertain stable behavior. The data analyzed are from the final 1–4 stimuli.

### Data analysis and statistics

The electrophysiological data of the TEVC measurements were analyzed with Ana (Dr. Michael Pusch, Genoa, Italy) and OriginPro 2024 (OriginLab Corporation, Northampton, MA, USA).

The significance of the mean differences was analyzed using one-way ANOVA with post-hoc mean comparison (Tukey test), where ns stands for not significant, p > 0.05, * for p ≤ 0.05, ** for p ≤ 0.01 and *** for p ≤ 0.001.

## Acknowledgement

We thank Anne Humberg and Thorsten König for excellent technical support.

## Supplementary material

**Suppl. Figure 1:**
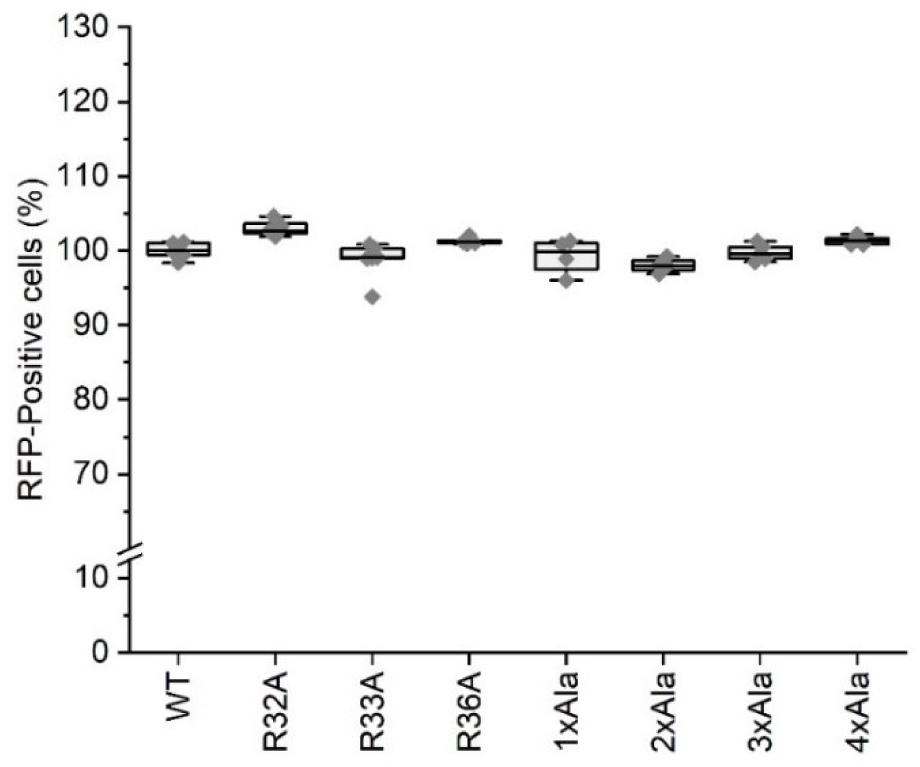
Similar expression of KCNE1 variants in HEK293 cells used for the *in vitro* cleavage assay. The red fluorescence of HEK293 cells transfected with red-fluorescently labeled KCNE1WT and KCNE1 variants was analyzed in FACS analyses and showed very similar expression levels.

**Suppl. Figure 2:**
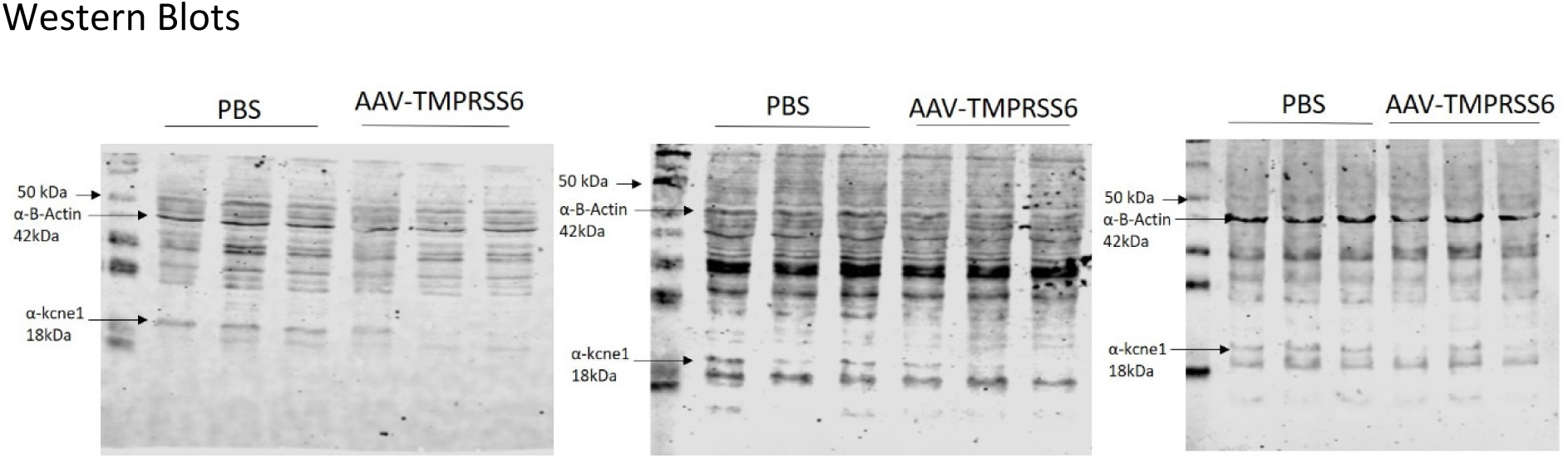
Western blots of Mouse hepatocytes show decreased abundance of KCNE1 in TMPRSS6-overexpressing cells.

**Suppl. Figure 3:**
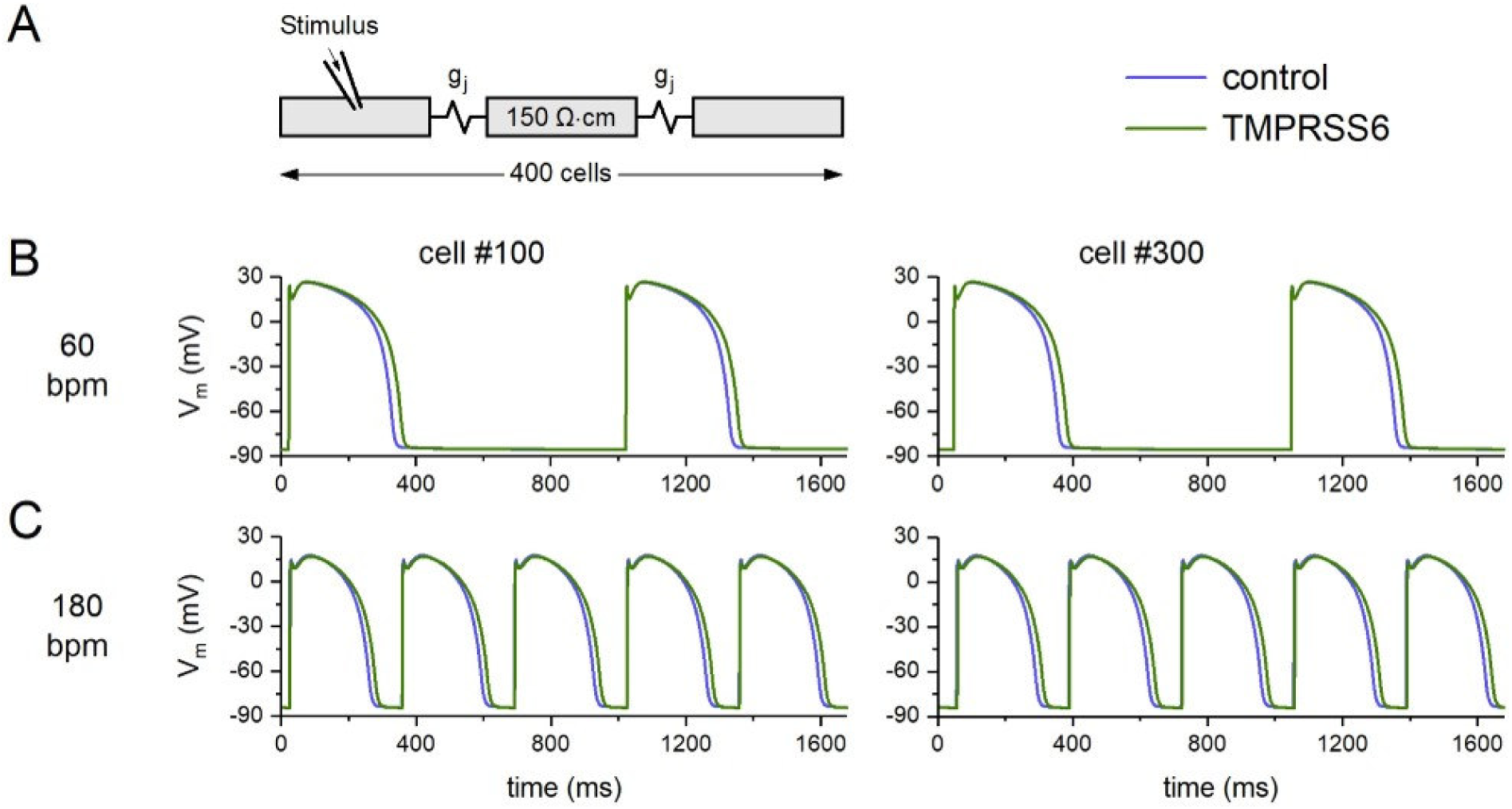
Effects of TMPRSS6 on the electrical activity in a simulated linear strand of human left ventricular cardiomyocytes under control conditions (blue traces) and during the TMPRSS6-induced reduction in I_Ks_ (green traces). **A;** Geometry of the unbranched strand of 400 cells. The gap junctional conductance (g_j_) was set to 10 µS and the myoplasmic resistivity was set to 150 Ω·cm. A suprathreshold stimulus was applied to the leftmost cell of the strand (cell #1). **B;** Membrane potential (V_m_) of cell #100 (left) and cell #300 (right) at a stimulus rate of 60 beats/min. **C;** V_m_ of cells #100 (left) and #300 (right) at a stimulus rate of 180 beats/min.

**Suppl. Figure 4:**
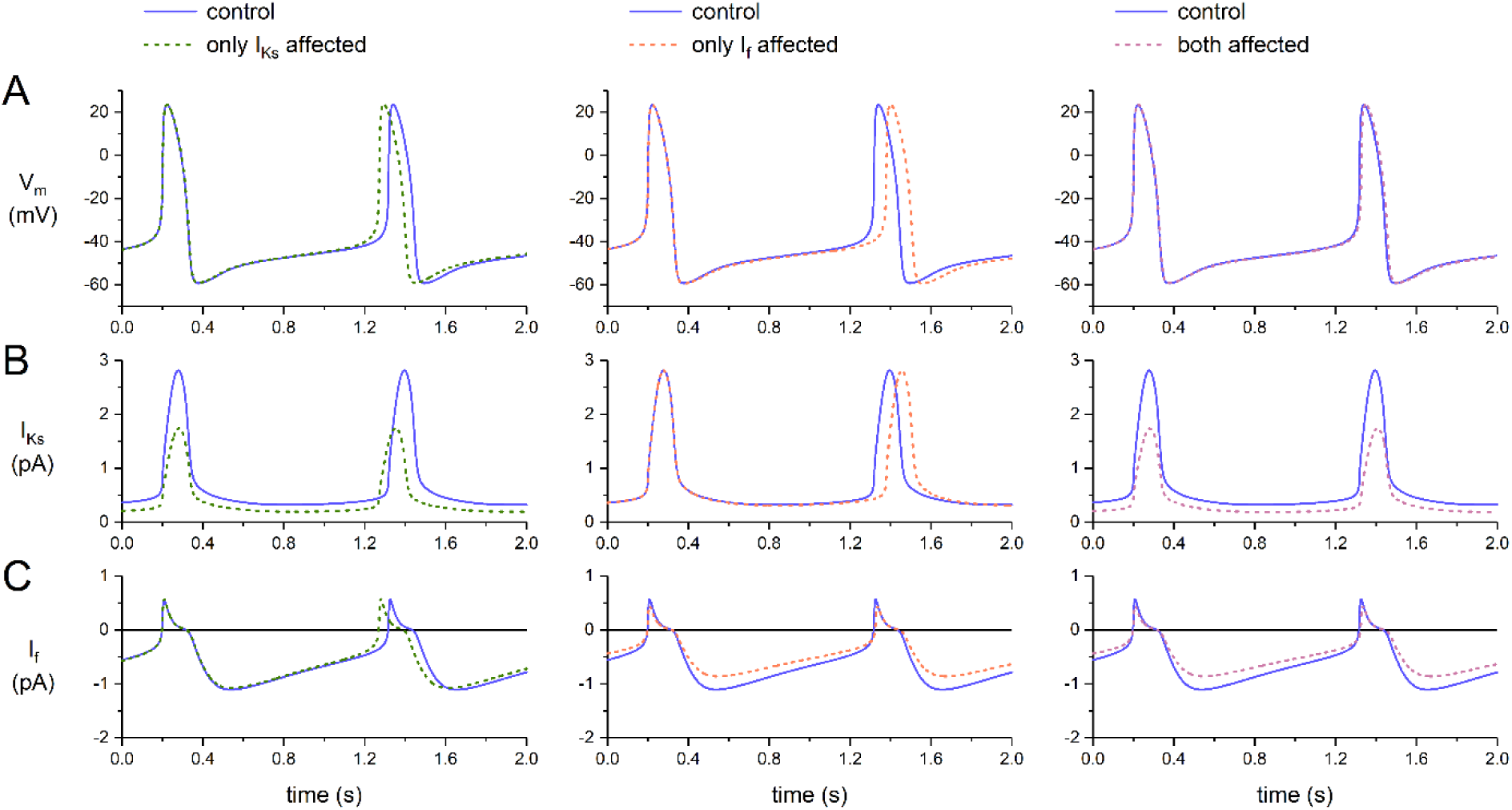
*In silico* effects of TMPRSS6 on the electrical activity of a single human sinus node cell under vagal tone, simulated by the presence of acetylcholine at a concentration of 10 μM. **A;** Membrane potential (V_m_), **B;** slow delayed rectifier potassium current (I_Ks_), and **C;** hyperpolarization-activated ‘funny’ current (I_f_) when TMPRSS6 affects I_Ks_ only (left), I_f_ only (middle), or both I_Ks_ and I_f_ (right).

**Suppl. Figure 5:**
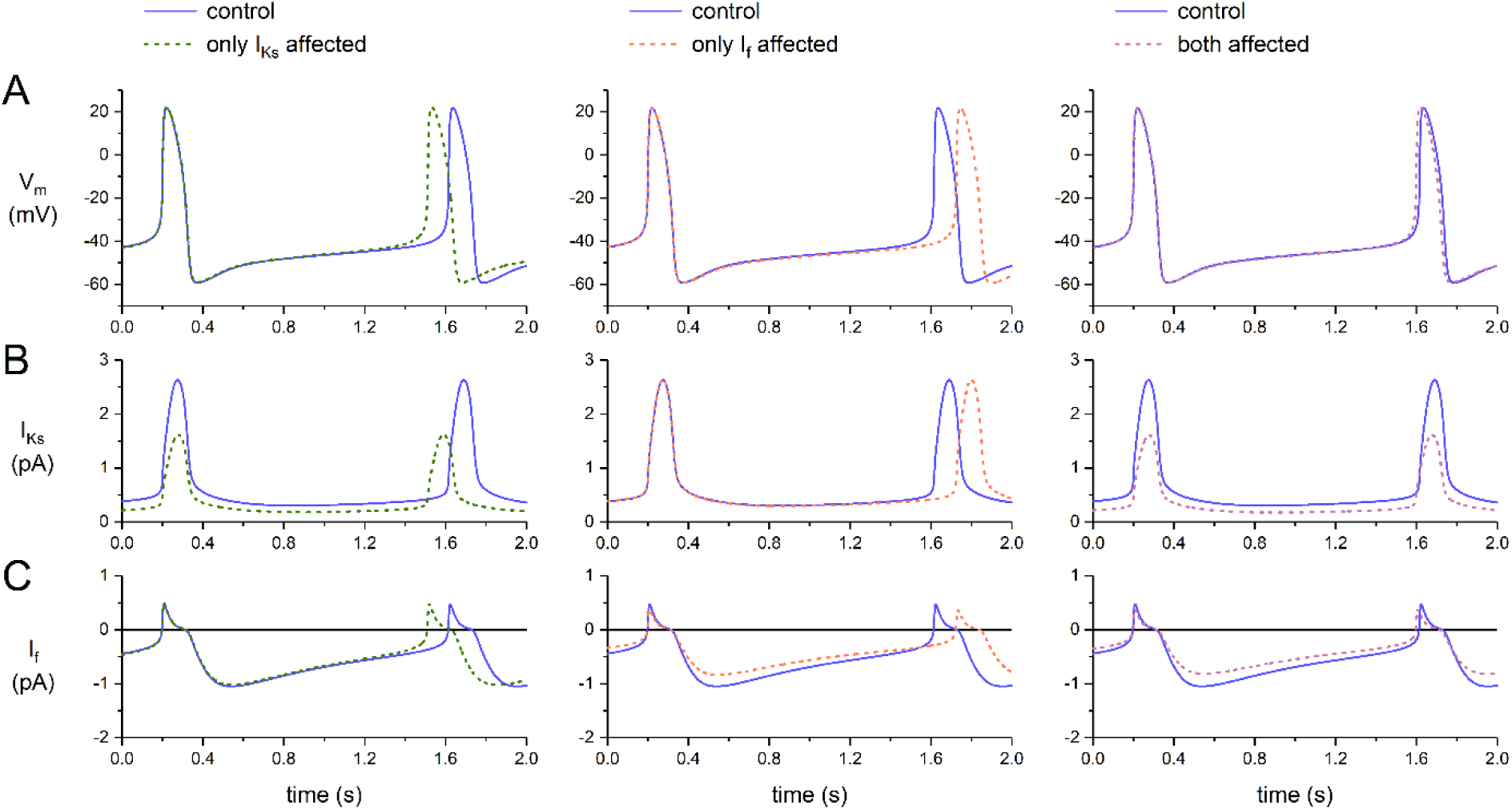
*In silico* effects of TMPRSS6 on the electrical activity of a single human sinus node cell under vagal tone, simulated by the presence of acetylcholine at a concentration of 20 μM. **A;** Membrane potential (V_m_), **B;** slow delayed rectifier potassium current (I_Ks_), and **C;** hyperpolarization-activated ‘funny’ current (I_f_) when TMPRSS6 affects I_Ks_ only (left), I_f_ only (middle), or both I_Ks_ and I_f_ (right).

**Suppl. Figure 6:**
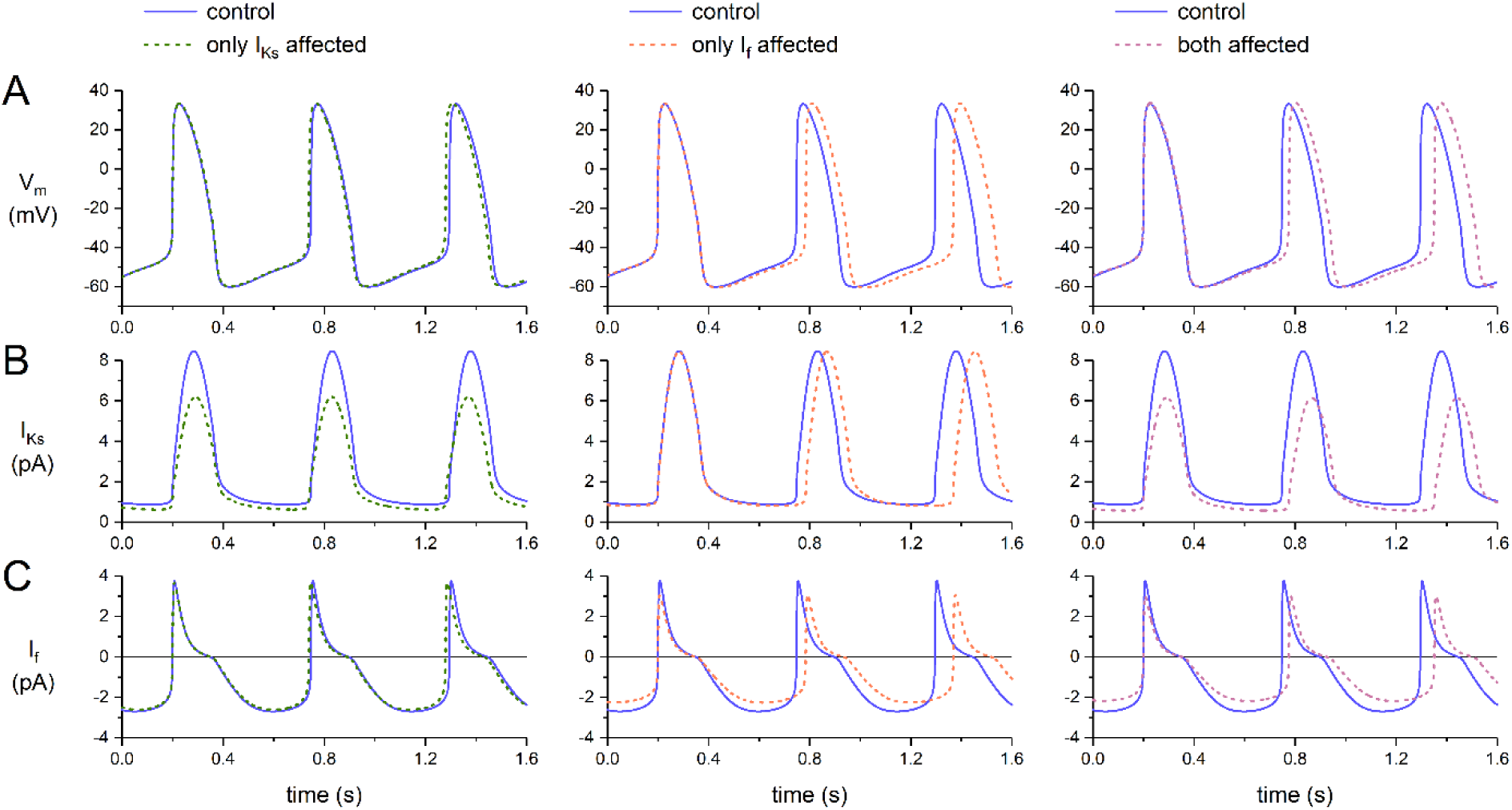
*In silico* effects of TMPRSS6 on the electrical activity of a single human sinus node cell under under β-adrenergic tone (‘high Iso’ settings ^49^). **A;** Membrane potential (V_m_), **B;** slow delayed rectifier potassium current (I_Ks_), and **C;** hyperpolarization-activated ‘funny’ current (I_f_) when TMPRSS6 affects I_Ks_ only (left), I_f_ only (middle), or both I_Ks_ and I_f_ (right). Note the differences in the ordinate and abscissa scales with Suppl. Figures 3 and 4.

**Suppl. Figure 7:**
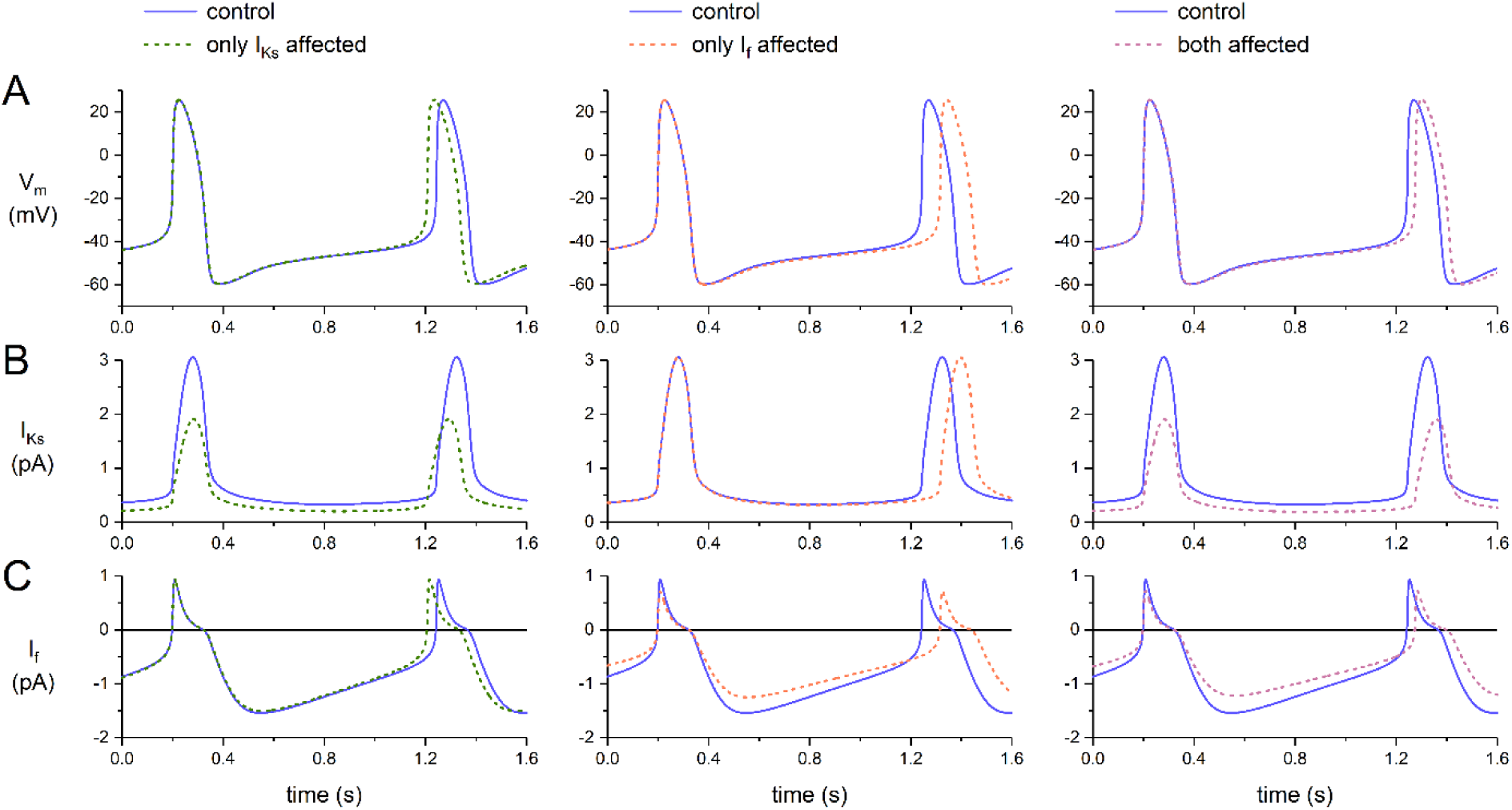
*In silico* effects of TMPRSS6 on the electrical activity of a single human sinus node cell in the presence of a passive atrial load of 0.4 pS/pF. **A;** Membrane potential (V_m_), **B;** slow delayed rectifier potassium current (I_Ks_), and **C;** hyperpolarization-activated ‘funny’ current (I_f_) when TMPRSS6 affects I_Ks_ only (left), I_f_ only (middle), or both I_Ks_ and I_f_ (right).

**Suppl. Figure 8:**
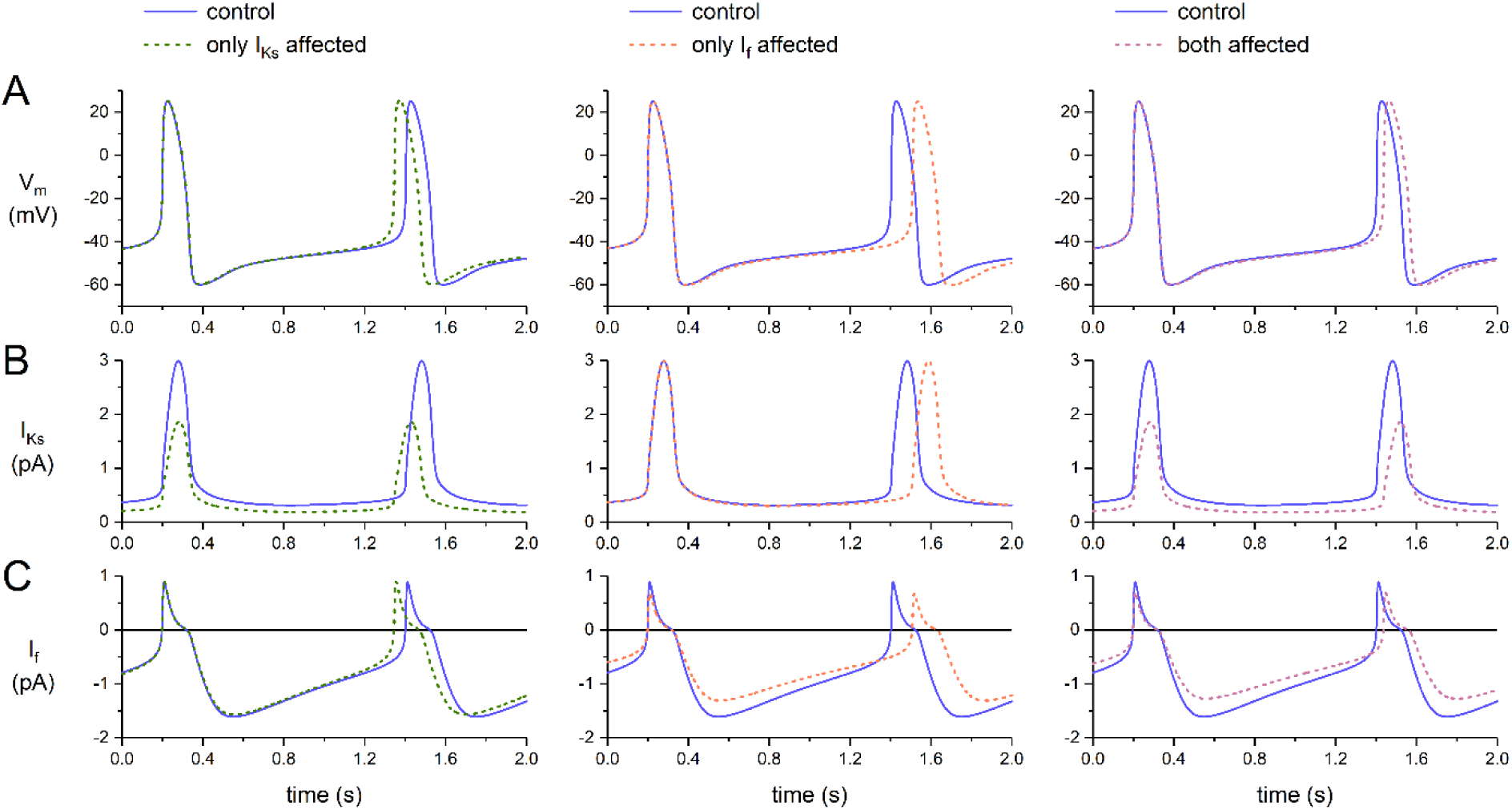
*In silico* effects of TMPRSS6 on the electrical activity of a single human sinus node cell in the presence of a passive atrial load of 0.6 pS/pF. **A;** Membrane potential (V_m_), **B;** slow delayed rectifier potassium current (I_Ks_), and **C;** hyperpolarization-activated ‘funny’ current (I_f_) when TMPRSS6 affects I_Ks_ only (left), I_f_ only (middle), or both I_Ks_ and I_f_ (right). Note the differences in the time scales with Suppl. Figure 6.

**Suppl. Figure 9:**
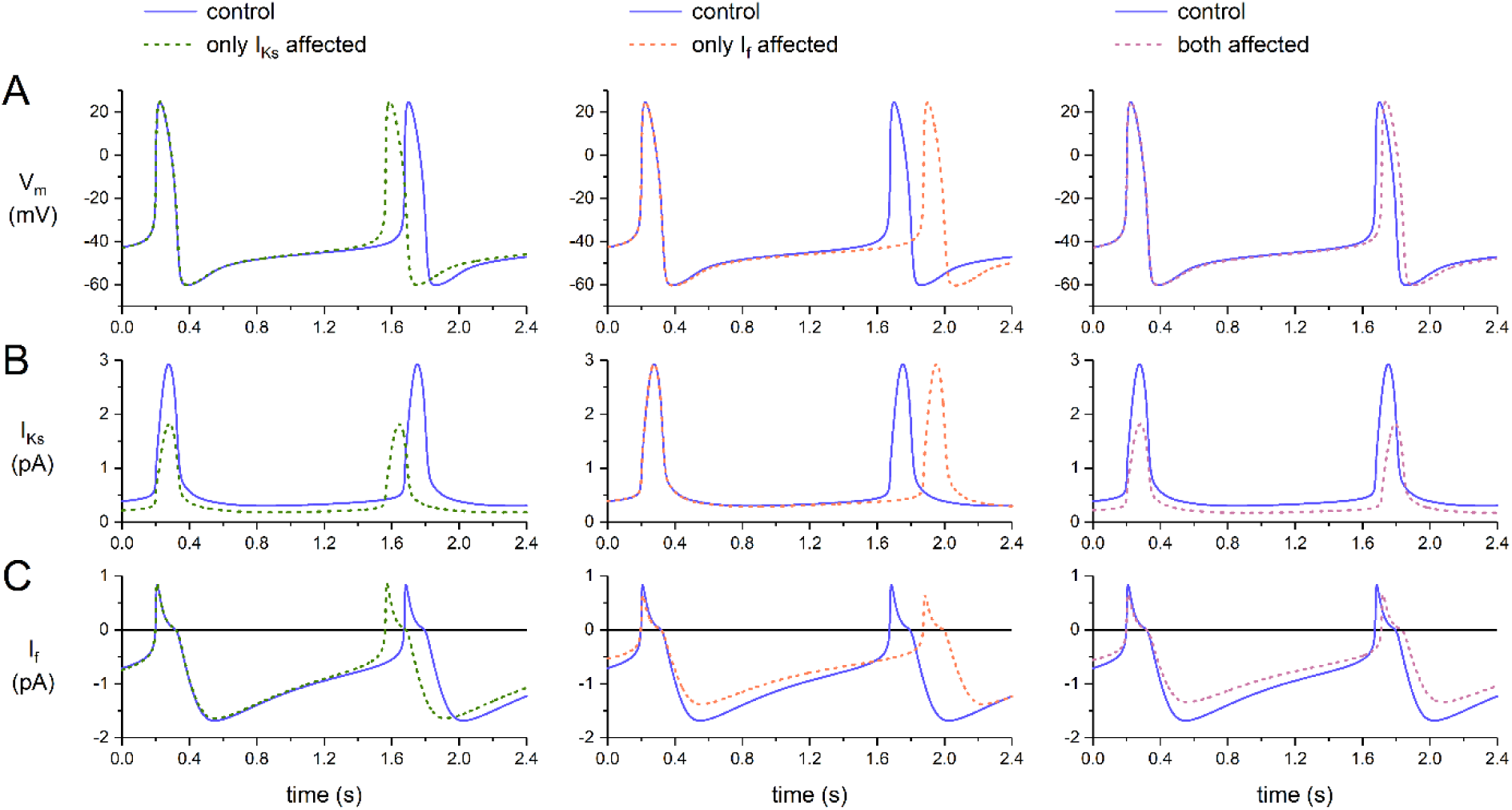
*In silico* effects of TMPRSS6 on the electrical activity of a single human sinus node cell in the presence of a passive atrial load of 0.8 pS/pF. **A;** Membrane potential (V_m_), **B;** slow delayed rectifier potassium current (I_Ks_), and **C;** hyperpolarization-activated ‘funny’ current (I_f_) when TMPRSS6 affects I_Ks_ only (left), I_f_ only (middle), or both I_Ks_ and I_f_ (right). Note the differences in the time scales with Suppl. Figures 6 and 7.

